# Nearest Neighbor Parameters for Estimating the Folding Stability of RNA Including Pseudouridine

**DOI:** 10.64898/2026.05.16.725682

**Authors:** Thandolwethu S. Shabangu, Elzbieta Kierzek, Sebastian Arteaga, Gregory S. Orf, Julia Stone, Olivia M. Hiltke, Megan Miaro, Elizabeth A. Jolley, Marta Soszyńska-Jóźwiak, Marta Szabat, Sharon Aviran, Philip C. Bevilacqua, Brent M. Znosko, Ryszard Kierzek, David H. Mathews

## Abstract

Nearest neighbor parameters are widely used in software for estimating the conformational stability of an RNA sequence folding into a specific structure. Folding stability for RNA with canonical nucleotides A, C, G, and U has been widely studied, but the same is not true for most modified nucleotides. In this work, we present a comprehensive set of nearest neighbor parameters for estimating the folding stability of RNA including pseudouridine in helical or loop contexts. These parameters are derived from 210 optical melting experiments involving helices with pseudouridine-A and pseudouridine-G pairs and with pseudouridine in loop motifs. The experiments include sequences with pseudouridine and U in the same strand, including U-A and U-G pairs, allowing us to consider the folding stability of sequences with both U and pseudouridine. On average, pseudouridine stabilizes RNA folding compared to U in an analogous motif, although this effect is sequence-context dependent. These parameters improve the modeling of folding stability for RNA secondary structures containing pseudouridine. We demonstrate that these parameters successfully model the secondary structure change for *Saccharomyces cerevisiae* U2 snRNA when two additional inducible pseudouridines are present. These parameters are freely available and incorporated into the RNAstructure software package.

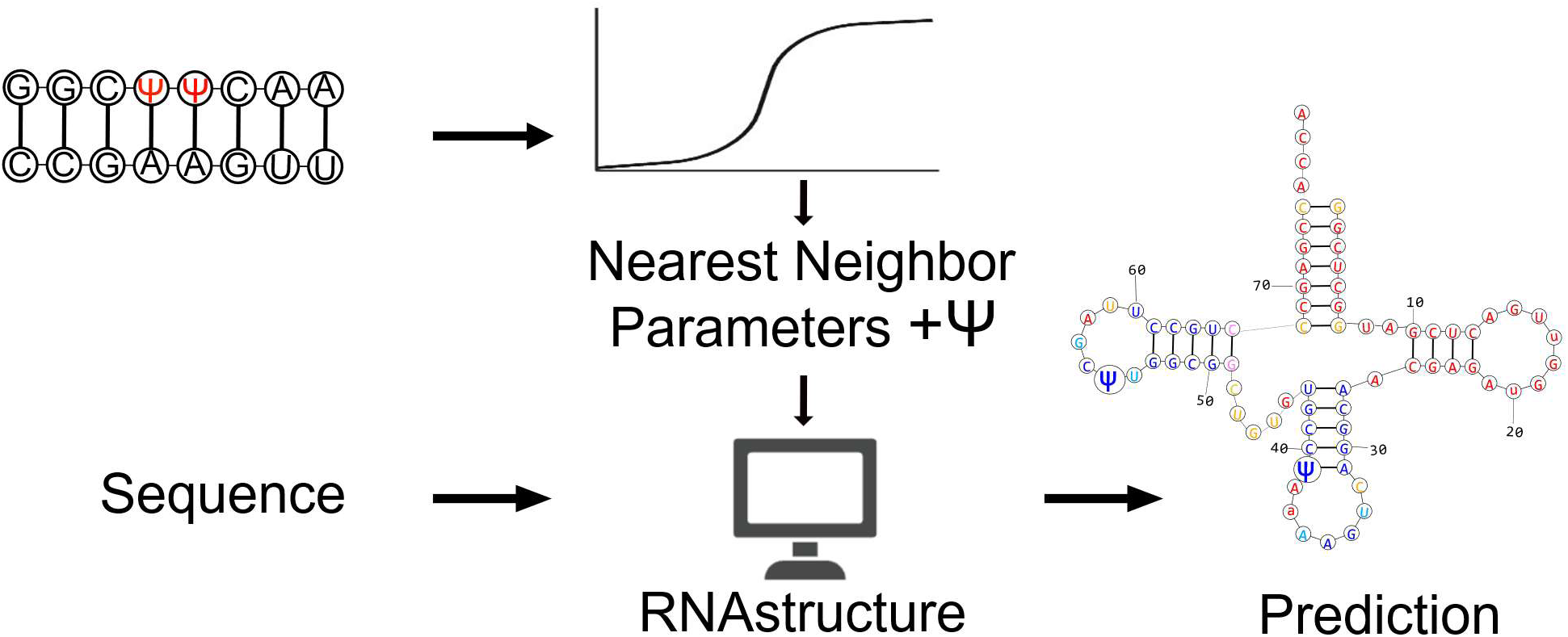

## Introduction

RNA plays diverse roles in biology, including regulating gene expression, catalyzing biochemical reactions, and acting as a mediator in protein synthesis [1–3]. The function of RNA is determined by its structure, which is organized into primary, secondary, and tertiary levels [4,5]. The secondary structure, which is defined by canonical base pairing, provides enough information to identify non-coding RNAs, develop a hypothesis about function, and design novel RNAs [6–14].

RNA modifications expand the RNA sequence repertoire by altering the canonical A, C, G, and U bases [15]. These modifications are numerous because covalent changes can occur in all four bases and the sugar [16,17]. They function as post-transcriptional regulators that affect folding stability, stability against degradation, structure, and RNA-protein interactions [18]. They participate in pre-mRNA splicing, nuclear export, and mRNA translation and contribute to immune cell functions that regulate immune responses [15,18,19].

Pseudouridine (Ψ) is the most abundant modified nucleotide and is found in various families of RNA across many species [20,21]. It is commonly found in positions where structure is important for function [21]. For example, mutants lacking specific Ψ residues in tRNA exhibit difficulties in translation, display slow growth rates, and fail to compete effectively with wild-type strains in mixed culture [21]. rRNAs containing Ψ exhibit subtle structural and stability differences compared to their U analogues, implying potential roles such as maintaining translation accuracy [22].

Pseudouridylation, the transformation of U to Ψ, occurs in two ways. The first is an RNA guide-independent mechanism, which involves stand-alone pseudouridylases, and the second is an RNA guide-dependent mechanism that is catalyzed by a family of box H/ACA ribonucleoproteins [20]. Pseudouridine differs from U by having a carbon–carbon glycosidic bond (Figure 1A), which allows greater rotational freedom of the base and introduces an additional hydrogen bond donor. These features give Ψ distinct chemical properties [20,21,23]. Pseudouridine makes the RNA backbone more rigid through a water-mediated hydrogen bond network and stabilizes base pairing and base stacking, which increases the folding stability of the RNA [20,24].

**Figure 1:**
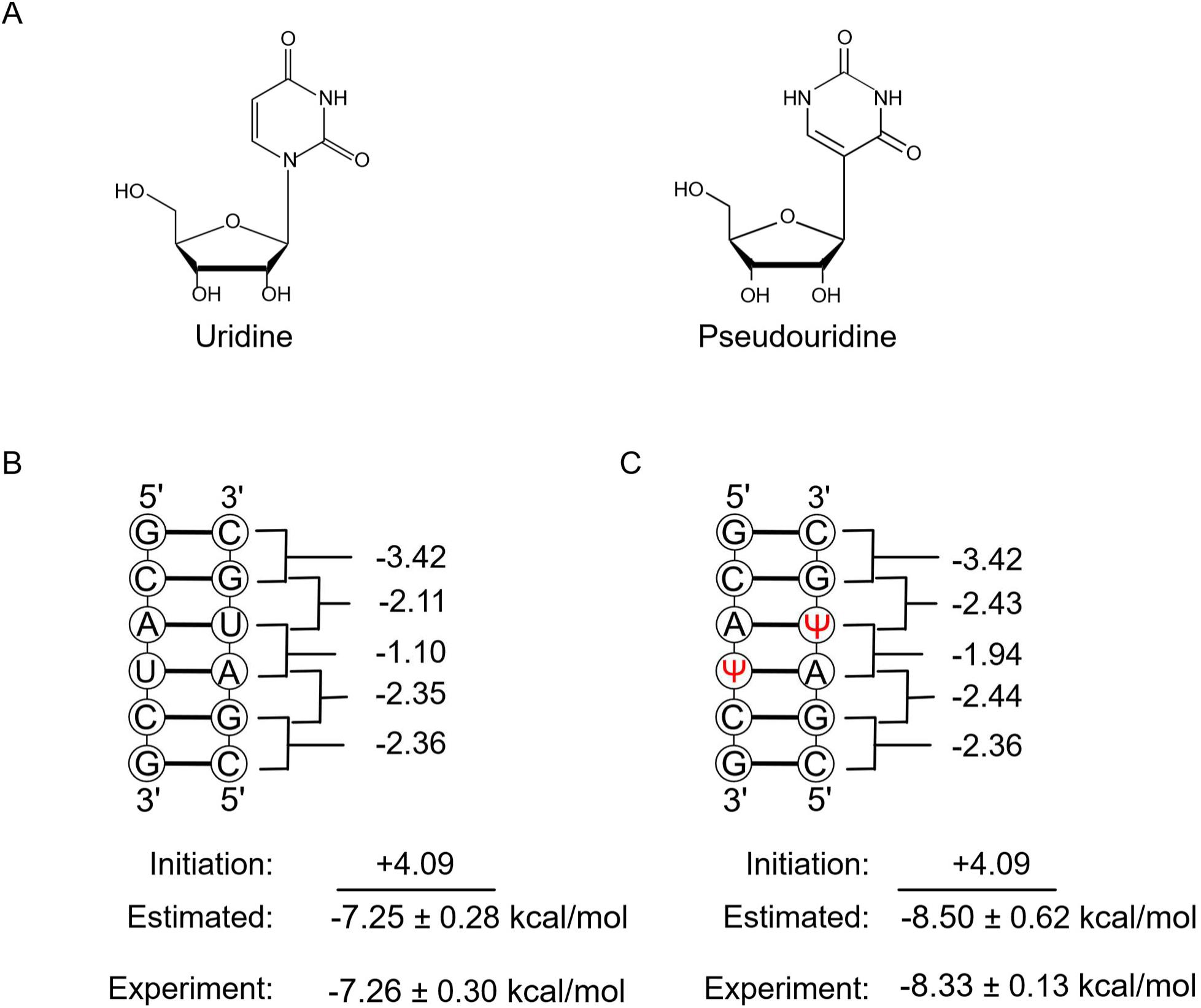
Pseudouridine stabilizes structure formation. Panel (**A**) compares uridine to pseudouridine. After pseudouridylation, the base N1 position becomes available for hydrogen bond donation. Panel (**B**) illustrates the stability calculation for 5’GCAUCG/3’CGUAGC and Panel (**C**) shows the stability calculation for 5’GCAΨCG/3’CGΨAGC. The calculations use parameters derived in this work and those reported previously [37,48]. For these examples, the total stability is obtained by summing the nearest neighbor stability increments and the initiation term. For self-complementary sequences, an additional symmetry correction is added. Additionally, for helices that terminate in A-U, A-Ψ, or G-Ψ ends, additional stability penalties are added. Uncertainty estimates are propagated from the uncertainty of the individual terms as explained in the Materials and Methods. This highlights the stabilizing effect that pseudouridine can confer on helices.

Nearest neighbor parameters can be used to estimate the free energy change of folding, i.e. the folding stability, to a secondary structure from a random coil [4,25]. The nearest neighbor parameters rely on two assumptions. First, the stability of a motif is determined by its sequence and the sequence of the adjacent base pairs. For helices, this is achieved using stacking terms of adjacent base pairs, where pairs at the ends of helices appear in one stack term, but pairs in the interior of helices appear in two stack terms. Second, the total folding free energy change of a secondary structure is the sum of the stabilities estimated for each motif [25]. An example calculation for a helix is shown in Figure 1B. These parameters are obtained through linear fits to folding stabilities, which are determined by optical melting experiments on small model structures [25]. They are widely used in RNA secondary structure prediction software [4,26–29]. Most algorithms use the Turner 2004 rules, which estimate the folding stability of RNA sequences with canonical base pairs [30] and were derived from 802 optical melting experiments [4,31,32].

Recently, parameters for predicting structures with m^6^A, A, C, G, and U were developed. These were derived from 45 optical melting experiments and included both helices and loop motifs [33]. Additional tests using 98 optical melting experiments confirmed that these parameters were as accurate for stability estimations with m^6^A as the A parameters are for estimations with sequences with A only [34]. RNAstructure, a software package for RNA secondary structure prediction and analysis, was updated to accept an alphabet of any size, including pairs that are orthogonal to the canonical base pairs, allowing the use of modified nucleotides [26,33,35].

In this work, we report parameters for predicting structures with Ψ, A, C, G, and U using the RNA Turner 2004 model as a basis [4,36], but with revised G-U pair parameters [37]. Previous work reported parameters for Ψ-A base pair helical stacks that were limited to single-substitution contexts, where only one U is replaced by Ψ within a stack or fully substituted stacks for which all U nucleotides are substituted with Ψ [10,38]. We report new helical stacking parameters for Ψ-A and Ψ-G pairs, adjacent to pairs with Ψ or canonical nucleotides including U, and parameters for Ψ-containing loop motifs. These parameters are based on a total of 210 optical melting experiments, of which 153 are new for this work. The choice of experiments was guided by a prior sensitivity analysis that identified the parameters with the most influence on the precision of secondary structure prediction [39].

The Ψ parameters determined herein are already publicly available in the RNAstructure software package, which improves modeling of RNA structures containing pseudouridine and enables estimation of their folding stability. To demonstrate the utility of these parameters, we modeled the conformational switch in U2 snRNA that is known to regulate splicing upon induction of two pseudouridines [40]. Our thermodynamic model recapitulates that the stem IIa structure predominates when three Ψs are present, whereas the stem IIc structure is favored with five Ψs.

## Materials and Methods

### Preparation of model RNA duplexes at The Institute for Bioorganic Chemistry

Oligonucleotides were synthesized on a BioAutomation MerMade12 DNA/RNA synthesizer using β-cyanoethyl phosphoramidite chemistry and commercially available phosphoramidites (ChemGenes, GenePharma, Glen Research), where standard protocols were followed as described previously [41,42]. For deprotection, oligoribonucleotides were treated with a mixture of 30% aqueous ammonia and ethanol (3:1 v/v) for 16 h at 55°C. Silyl protecting groups were removed with the use of triethylamine trihydrofluoride. The deprotected oligonucleotides were purified using silica gel thin layer chromatography in a mixture of 1-propanol, aqueous ammonia, and water (55:35:10 v/v/v), as described previously [43]. Mass spectrometry analyses (MALDI) was performed for most of the oligonucleotides.

### Preparation of model RNA duplexes at Saint Louis University

All oligonucleotides were ordered from Integrated DNA Technologies, Inc. (Coralville, IA) if no modified residues were in the sequence or Dharmacon (now Horizon Discovery Ltd.) if the sequence contained a pseudouridine residue. Although coupling efficiencies during the synthesis process were high, failure sequences were removed for all oligonucleotides. Standard procedures were followed [44,45]. The purity and identity of each oligonucleotide were verified on a Shimadzu liquid chromatography mass spectrometry (LCMS)-2010EV via manual injection or with a Prominence-i LC-2030 auto-sampler. Concentrations of oligonucleotides were calculated at 80 °C with recorded absorbances at 260 nm in a 0.1 cm cuvette using RNACalc [46,47] and the Beer-Lambert Law.

### UV melting experiments at The Institute for Bioorganic Chemistry

The thermodynamic measurements were performed for nine various concentrations of RNA duplex in the range 100 µM - 1 µM on JASCO V-650 UV/Vis spectrophotometer in buffer containing 1 M sodium chloride, 20 mM sodium cacodylate, and 0.5 mM Na_2_EDTA, pH 7.0 [46,48]. Oligonucleotide single strand concentrations were calculated from the absorbance above 80 °C, and single strand extinction coefficients were approximated by a nearest-neighbor model [49,50]. Absorbance vs. temperature melting curves were measured at 260 nm with a heating rate of 1 °C/min from 4 to 90 °C on a JASCO V-650 spectrophotometer with a thermoprogrammer. The temperatures of the measurements were recalibrated using a temperature calibration curve measured with a thermocouple and a program that uses a linear extrapolation [51]. The melting curves were analyzed, and the thermodynamic parameters were calculated from a two-state model with the program MeltWin 3.5 [52]. For most duplexes, the ΔH° parameters derived from T_M_^-1^ vs. log(C_T_/4) plots are within 15% of that derived from averaging the fits to individual melting curves, as expected if the two-state model is reasonable.

### UV melting experiments at Saint Louis University

The buffer conditions for this study were 20.0 mM sodium cacodylate with 1.0 M NaCl and 5.0 mM Na_2_EDTA. The solution was prepared at 23 °C and adjusted to a pH of 7.0. The absorbances were taken on a Beckman-Coulter DU800 spectrophotometer with an automated high-performance temperature controller. Absorbances were recorded at 260 nm while temperature increased at a rate of 1°C/minute from 10-90 °C across a 50-fold oligonucleotide concentration range. The plotted absorbance versus temperature graphs were then fit to extract thermodynamic parameters.

The MeltR [53] software was used to fit all optical melting curves. Raw data was converted to Tidy format and fit using MeltR as an R-markdown script. The meltR.A function was used to acquire initial thermodynamic parameters with no baseline trimming of the upper and lower baselines of the melt curves. The meltR.A data was saved to a variable input into the BLTrimmer function for automated baseline trimming of upper and lower baselines. Statistically driven upper and lower baseline confidence intervals were extracted with accompanying thermodynamic parameters. The van’t Hoff analysis data was used in the nearest-neighbor derivations.

### Reproducibility

To test the reproducibility of optical melting experiments across labs, experiments were performed at Saint Louis University, St. Louis, and at the Institute for Bioorganic Chemistry, Poznan, on three calibration duplexes that had previously been studied at the University of Rochester [48,54,55]. These sequences are (CGCGCG)_2_, 5’GCUACG/3’CGAUGC, and (GCGUUCGC)_2_. The results were recently reported and are summarized here in Supplementary Table S1 [51]. The largest difference between sites is 0.32 kcal/mol for 5’GCUACG/3’CGAUGC, which represents 4.2% of the mean ΔG°_37_ of 7.62 kcal/mol. This is consistent with prior studies of reproducibility across sites [48,56]. Here it was important to demonstrate reproducibility with prior work because the new pseudouridine parameters are integrated with the Turner 2004 parameters. Also, the parameters are fit using experiments across two sites, Saint Louis University and the Institute for Bioorganic Chemistry.

### Helical stacking parameters

To determine the RNA helix stacking nearest neighbor parameters, we performed non-error-weighted linear regression using the Statsmodels [57] library in Python. We used duplexes with canonical pairs, including Ψ-A and Ψ-G pairs, shown in Supplementary Table S2, along with 24 duplexes reported by Hudson et al. [38] and one duplex reported by Hall et al. [58].

We first assessed if each duplex denaturation was consistent with two state behavior by calculating the percent difference in ΔH° values fit by two methods, the average of curve fits and T_M_^-1^ vs log C_T_ fits. Agreement within 15% is considered consistent with two-state denaturation [46,59,60]. Seven duplexes that had differences of greater than 15% were therefore excluded from the fit (identified in Table S2).

We used the ΔG°_37_ values from T_M_^-1^ vs log C_T_ plots method to fit the stacks. For each duplex, to isolate the stability contribution by pairs with Ψ, we subtracted the duplex initiation term, the terminal A-U penalty term (for duplexes ending in A-U), the symmetry term (for self-complementary duplexes), stack terms for Watson-Crick-Franklin pairs [48], and stack terms for G-U pairs [37]. The standard error of the regression was used as the estimate of uncertainty.

A plot of the residuals of the duplexes used in the stacks (ΔG°_experiement_ – ΔG°_predicted_) revealed one outlier experiment, 5’GCUΨCGC/3’CGAGGCG (Supplementary Figure S1A). The data was then refitted with this experiment excluded. Supplementary Figure S1B shows the residual plot after this sequence was excluded. Supplementary Table S3 shows the stacking parameters for Ψ-A pairs and the number of occurrences of each stacking parameter in the set of helices. Supplementary Table S4 shows the Ψ-G stack parameters.

### Dangling ends

We studied 12 duplexes to determine the stabilities of dangling end motifs, which included Ψ as both the dangling end, the terminal base pair, or both (Supplementary Table S5). A subset of ten duplexes were self-complementary, resulting in two dangling ends, and two were non-self-complementary, resulting in one dangling end. We used the free energy change values from the T_M_^-1^ vs log C_T_ plots fit of the optical melting experiments.

The ΔG°_37_ for each motif was calculated (Supplementary Table S6) as shown below:

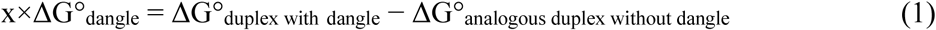

where the value of x is 2 for self-complementary duplexes with two identical dangling end motifs. For non-self-complementary sequences, x is 1. The ΔG°_duplex with dangle_ values are those that were measured from the optical melting experiments. The ΔG°_analogous duplex without dangle_ are free energy changes for duplexes with identical sequence but without the dangling end. The experimental values were used for ΔG°_analogous duplex without dangle_ when available, otherwise the stability was estimated using the helical stack parameters.

In three cases, the same dangling end was measured multiple times, and the mean value is used for the nearest neighbor parameters. These cases are the 5’ dangling Ψ on a U-A terminal pair, the 3’ dangling Ψ on a U-A terminal pair, and the 3’ dangling Ψ in a G-C terminal pair. In all cases, the agreement was good with standard errors of the mean of 0.025, 0.1625, and 0.0575 kcal/mol, respectively. The duplex (AUGCAΨΨ)_2_ has a dangling end that was apparently destabilizing. Because we expect dangling ends to stabilize helix formation, we excluded this sequence from further analysis. We assume this apparent destabilization is an artifact caused by uncertainty in the measurements.

To extrapolate the values for the dangling ends that were not experimentally studied, we calculated:

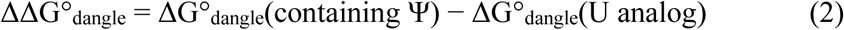

The ΔG°_dangle_(U analog) is the value of the dangle for the analogous dangling end that has U replacing Ψ, as tabulated in [39].

The mean ΔΔG°_dangle_ for the Ψ 5’ dangling ends that were measured and the mean ΔΔG°_dangle_ for the measured Ψ 3’ dangling ends were -0.44 ± 0.07 kcal/mol and -0.34 ± 0.06 kcal/mol, respectively. To extrapolate to dangling Ψ values we did not measure, we added the corresponding mean ΔΔG°_dangle,_ (5’ or 3’) to the analogous U dangling end.

To estimate the additional stability for 5’ dangling A, C, G, and U on terminal A-Ψ or terminal Ψ-A pairs, we used the measured ΔΔG°_dangle_ of dangling C on an A-Ψ terminal pair (−0.26 ± 0.39 kcal/mol). We added this ΔΔG°_dangle_ to the analogous U values to obtain the corresponding ΔG°_dangle_.

To estimate the additional stability for 3’ dangling A, C, G and U on a terminal A-Ψ pair, we used the measured ΔΔG°_dangle_ value of the dangling A (−0.26 ± 0.51 kcal/mol). For 3’ dangling A, C, G and U on a Ψ-A pair, we used the measured ΔΔG°_dangle_ of the dangling G (+0.67 ± 0.46 kcal/mol). We added these ΔΔG°_dangle_ values to analogous U values to obtain the corresponding ΔG°_dangle_ for these motifs. Interestingly, this resulted in the 3’ dangle of C and U on the Ψ-A having positive values of 0.54 ± 0.47 and 0.58 ± 0.49 kcal/mol, respectively. Although we expect dangling ends to be stabilizing, we kept these values.

In the Turner 2004 RNA model, dangling ends on G-U and U-G pairs are set equal to the equivalent dangle on an A-U or U-A pair [32]. Following this model, we set dangling end values for G-Ψ and Ψ-G terminal pairs equal to dangling ends on A-Ψ and Ψ-A terminal pairs, respectively.

### Terminal mismatches

14 optical melting experiments were used to derive the stabilities of terminal mismatch motifs, shown in Supplementary Table S7, which included Ψ as the terminal mismatch or the terminal base pair. We used the T_M_^-1^ vs log C_T_ plots fit of the optical melting experiments to calculate the motif stabilities (Supplementary Table S8) as shown below:

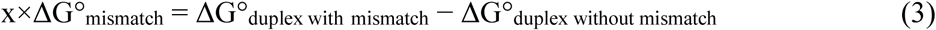

where x is 2 for self-complementary duplexes to account for the fact that the motif is present twice. For non-self-complementary duplexes, x is set to 1. The same terminal mismatch was measured twice for C-G followed by Ψ-Ψ, the mean value (−2.06 ± 0.17 kcal/mol) is used for the nearest neighbor parameters. The agreement was good between the two cases with a standard error of the mean of 0.12 kcal/mol. The sequence (AAUGCAΨC)_2_ was excluded from the model because the terminal mismatch was apparently destabilizing (Supplementary Table S8), which we assume is an artifact caused by uncertainty in measurements.

To extrapolate the stabilities of motifs that were not measured, we first calculated:

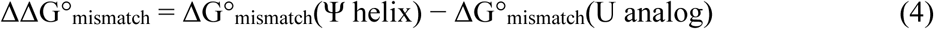

where ΔΔG°_mismatch_ is the difference in stability between a mismatch containing Ψ and a mismatch with U in the analogous position, as tabulated by [39,61].

To extrapolate the values for a C-Ψ following an A-U, C-G or U-A pair, we added the measured ΔΔG°_mismatch_ of a G-C next to a C-Ψ terminal pair (−0.31 ± 0.62 kcal/mol) to the analogous C-U terminal mismatch values for these pairs.

To estimate the value of U-Ψ next to an A-U, C-G, or U-A pair, we added the measured ΔΔG°_mismatch_ of G-C next to a U-Ψ pair (−0.64 ± 0.36 kcal/mol) to the analogous U-U terminal mismatch values for these pairs. For Ψ-C following an A-U, C-G and U-A pair, we added the measured ΔΔG°_mismatch_ value of G-C next to Ψ-C (−0.54 ± 0.63 kcal/mol) to the analogous U-C terminal mismatch values of these pairs. To estimate the stabilities of Ψ-U next to A-U, C-G and U-A, we added the measured ΔΔG°_mismatch_ value of G-C next to Ψ-U (−1.06 ± 0.64 kcal/mol) to the analogous U-U terminal mismatch values of these pairs.

To extrapolate the value of Ψ-Ψ next to U-A, we added the mean ΔΔG°_mismatch_ value of Ψ-Ψ next to A-U, G-C, and C-G (−0.57 ± 0.20 kcal/mol) to the analogous U-U value for this mismatch.

The mean ΔΔG°_mismatch_ of A-A and C-A next to A-Ψ (−0.56 ± 0.23 kcal/mol) was added to the analogous A-C, A-G, C-C, C-U, G-A, G-G, U-C, and U-U next to A-Ψ to extrapolate the stabilities for these terminal mismatches. The ΔΔG°_mismatch_ of A-Ψ next to Ψ-Ψ (−0.51 ± 0.52 kcal/mol) was added to the analogous U-U values of A-Ψ next to C-Ψ, U-Ψ, Ψ-C, and Ψ-U to extrapolate the stabilities of these terminal mismatches. The ΔΔG°_mismatch_ of the measured Ψ-A next to A-C (−0.68 ± 0.68 kcal/mol) was added to the analogous A-A, A-G, G-A, C-C, C-U, G-G, U-C, and U-U next to U-A to extrapolate the values of these terminal mismatches.

To estimate the stability of Ψ-A next to C-Ψ, the sum of ΔΔG°_mismatch_ of C-Ψ next to G-C and A-C next to Ψ-A (+0.37 ± 0.93 kcal/mol) was added to the analogous C-U next to U-A stability. To estimate the stability of Ψ-A next to U-Ψ, the sum of ΔΔG°_mismatch_ of Ψ-A next to A-C and G-C next to U-Ψ (+0.04 ± 0.78 kcal/mol) was added to the analogous U value of U-A next to U-U. To estimate the stability of Ψ-A next to Ψ-C, the sum of ΔΔG°_mismatch_ of Ψ-A next to A-C and G-C next to Ψ-C (+0.14 ± 0.94 kcal/mol) was added to the analogous U value of U-A next to U-C.

To estimate the stability of Ψ-A next to Ψ-U, the sum of ΔΔG°_mismatch_ of Ψ-A next to A-C and ΔΔG°_mismatch_ of G-C next to Ψ-U (−0.39 ± 0.94 kcal/mol) was added to the analogous U-A next to U-U. To estimate the stability of Ψ-A next to Ψ-Ψ, the sum of ΔΔG°_mismatch_ of Ψ-A next to A-C and the mean of ΔΔG°_mismatch_ of Ψ-Ψ next to A-U, G-C and C-G (+0.10 ± 0.72 kcal/mol) was added to the analogous value of U-A next to U-U.

Following the Turner 2004 model, in which the stability of an analogous mismatch on an A-U or U-A pair was used to estimate mismatches on G-U or U-G pairs when there was no experiment [62], we set the mismatches on Ψ-G and G-Ψ terminal mismatches equal to the mismatches on the analogous Ψ-A and A-Ψ terminal pairs, respectively.

### Internal loops/Bulge loops

The duplexes used to determine the stabilities of internal loops and bulge loops are shown in Supplementary Table S9 and Supplementary Table S10, respectively. The free energy changes from T_M_^-1^ vs log C_T_ plots fit of the optical melting experiements were used for the fit. The stabilities of the motifs were determined as follows:

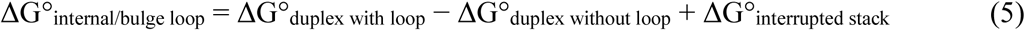

where ΔG°_interrupted stack_ is the stability of the helical stack parameter for the stack that is present in the duplex without the loop, but is interrupted by the loop in the duplex with the loop [63]. An additional U-A, Ψ-A, and Ψ-G terminal pair penalty of 0.45 ± 0.04 kcal/mol, 0.42 ± 0.21 kcal/mol and 0.21 ± 0.15, respectively, is applied to ΔG°_interrupted stack_ each time such a pair is present in the stack. The one exception to Equation 5 is that single nucleotides bulges are assumed to not interrupt the helical stack. Therefore, ΔG°_interrupted stack_ is set to 0 kcal/mol for single nucleotide bulge loops.

When an experimental measurement for the duplex without the loop is not available, we subtract the sum of the helical stacking parameters, the terminal pair penalties for A-U, Ψ-A, or Ψ-G (applied to each terminal pair), and the intermolecular initiation term from ΔG°_duplex with loop_.

The motif stabilities for bulge loops and internal loops are shown in Supplementary Table S11 and Supplementary Table S12, respectively. The bulge loop in the sequence 5’GCGΨGCG/3’CGC-CGC had unusually high stability (+0.09 ± 0.59 kcal/mol) compared to other buldge loops, so it was excluded from the estimates of the nearest neighbor parameters.

In the Turner 2004 model, the internal loop stabilities are modeled using sequence-dependent parameters that account for the loop size, symmetry, and identity of the first mismatch [4]. To model the internal loop stabilities for the Ψ parameters, we use the Turner 2004 model and added Ψ-Ψ, Ψ-U, U-Ψ, and Ψ-C terms and an A-Ψ closure term. These first mismatches involving Ψ stabilize internal loop formation compared to analogous U-U first mismatches. We first calculated:

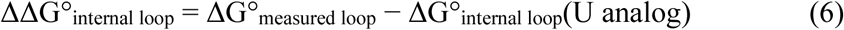

where ΔG°_measured loop_ is the stability calculated as described above. ΔG°_internal loop_(U analog) are the values when the internal loops have U instead of Ψ, either as reported for an experiment in [39] or as calculated as explained in [4]. The average of the ΔΔG°_internal loop_ values for duplexes with Ψ-Ψ first mismatch pairs was used to determine the Ψ-Ψ term (−1.68 ± 0.35 kcal/mol). The average of the ΔΔG°_internal loop_ values for Ψ-U and U-Ψ first mismatch pairs (−1.00 ± 0.34 kcal/mol) was used to calculate the Ψ-U and U-Ψ terms. The ΔΔG°_internal loop_ of the Ψ-C was measured for one sequence so we used this value for the Ψ-C term (−0.68 ± 0.61 kcal/mol). The average of the ΔΔG°_internal loop_ values for motifs with A-Ψ closures (−0.48 ± 0.10 kcal/mol) was used to calculate the A- Ψ closure term.

Following the Turner 2004 rules, for 1×1, 1×2, and 2×2 nucleotide loops that were experimentally measured, we used the experimental values. The sequence CΨGCΨGG_2_ was excluded from the model because it appears to be too stable in relation to other 1×1 nucleotide internal loops. It has ΔΔG°_internal loop_ of -4.84 ± 1.38 kcal/mol, substantially lower than all other measured internal loops. The Ψ-C term was applied exclusively to 1×1 nucleotide internal loops because only a single experimental measurement was available, and that measurement was a 1×1 nucleotide internal loop. This avoided overfitting the model. For 1×2 nucleotide internal loops, when we had to choose between applying the Ψ-U or Ψ-Ψ term, we used Ψ-Ψ because it provides the larger bonus. For 2×2 nucleotide internal loops, we added the bonuses for both the mismatches if they appear. For example, if an internal loop contains a Ψ-U base pair followed by a Ψ-Ψ base pair, we add both ΔΔG°_37_ terms.

### Hairpin loops

Hairpin loop stabilities were determined from 12 optical melting experiments (Supplementary Table S13). To calculate the stabilities from the measured optical melting experiments, we subtracted the stability of the helical stacks (estimated with nearest neighbor parameters but excluding the intermolecular initiation term).

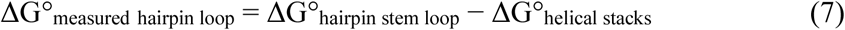

where ΔG°_hairpin stem loop_ is the experimentally determined stability of the hairpin with its closing stem. These results are shown in Supplementary Table S14, which also shows ΔG°_estimated_. The ΔG°_estimated._ values were calculated with the sum of a terminal mismatch, length-dependent initiation term, and sequence-dependent terms as described in the Nearest Neighbor Database https://rna.urmc.rochester.edu/NNDB/ [36].

### Propagation of uncertainties

We estimate the uncertainties under the assumption that the terms are uncorrelated and propagate them using:

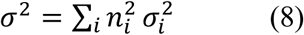

where σ_i_ is the uncertainty of the i^th^ parameter or experiment and n_i_ is the number of occurrences of that parameter or experiment [64]. For mean values, the uncertainty is reported as the standard error of the mean.

We estimate uncertainty in optical melting experiments as 4% of ΔG°_37_. This estimate follows Xia et al. [48] and is based on the reproducibility of optical melting experiments across sites.

### U2snRNA Structure Predictions

We downloaded the U2 snRNA sequence from the yeast snoRNA database, available at https://people.biochem.umass.edu/sfournier/fournierlab/snornadb/mastertable.php [65]. This catalogues nucleotide modifications, especially those created by small nucleolar RNA:protein complexes (SnoRNPs). U2 snRNA has three constitutively expressed pseudouridines at positions 35, 42, and 44, as well as two inducible pseudouridines at positions 56 and 93 [40]. We predicted the secondary structures and base pairing probabilities of U2 snRNA with three pseudouridines and with five pseudouridines using the Ψ alphabet of RNAstructure. We used RNAstructure *Probscan* [66] to quantify the probabilities of forming stem IIa or stem IIc structures in U2 snRNA.

## Results

### Helical stacking parameters

To fit helical stack parameters, we used 147 helices containing Ψ-A and Ψ-G base pairs in addition to the standard canonical pairs (Supplementary Table S2). Of those that were used in the linear regression fit, 110 were synthesized and melted for this study, 12 were previously studied in [67], 24 were reported in [38] and one was reported in [58].

Figure 2 compares the Ψ-A stacks to the analogous Watson-Crick-Franklin stack values for U. On average, substituting U with Ψ results in stabilization, with Ψ-A stacks being more stabilizing than U-A stacks by an average of -0.56 ± 0.13 kcal/mol. This effect is sequence-context dependent. The ΔΔG°_37,_ the difference between the Ψ-A stacks and their U analogues, ranges from -1.29 ± 0.34 kcal/mol to 0.01 ± 0.19 kcal/mol (Supplementary Table S3). This indicates a complete nearest neighbor model is required to estimate the stabilities of duplexes with Ψ-A and Ψ-G pairs, rather than relying on a single average value. Notably, some stacks exhibit comparable stability to their analogous U stacks, especially when considering the uncertainties, such as Ψ-A followed by Ψ-A (−0.92 ± 0.19 kcal/mol).

**Figure 2:**
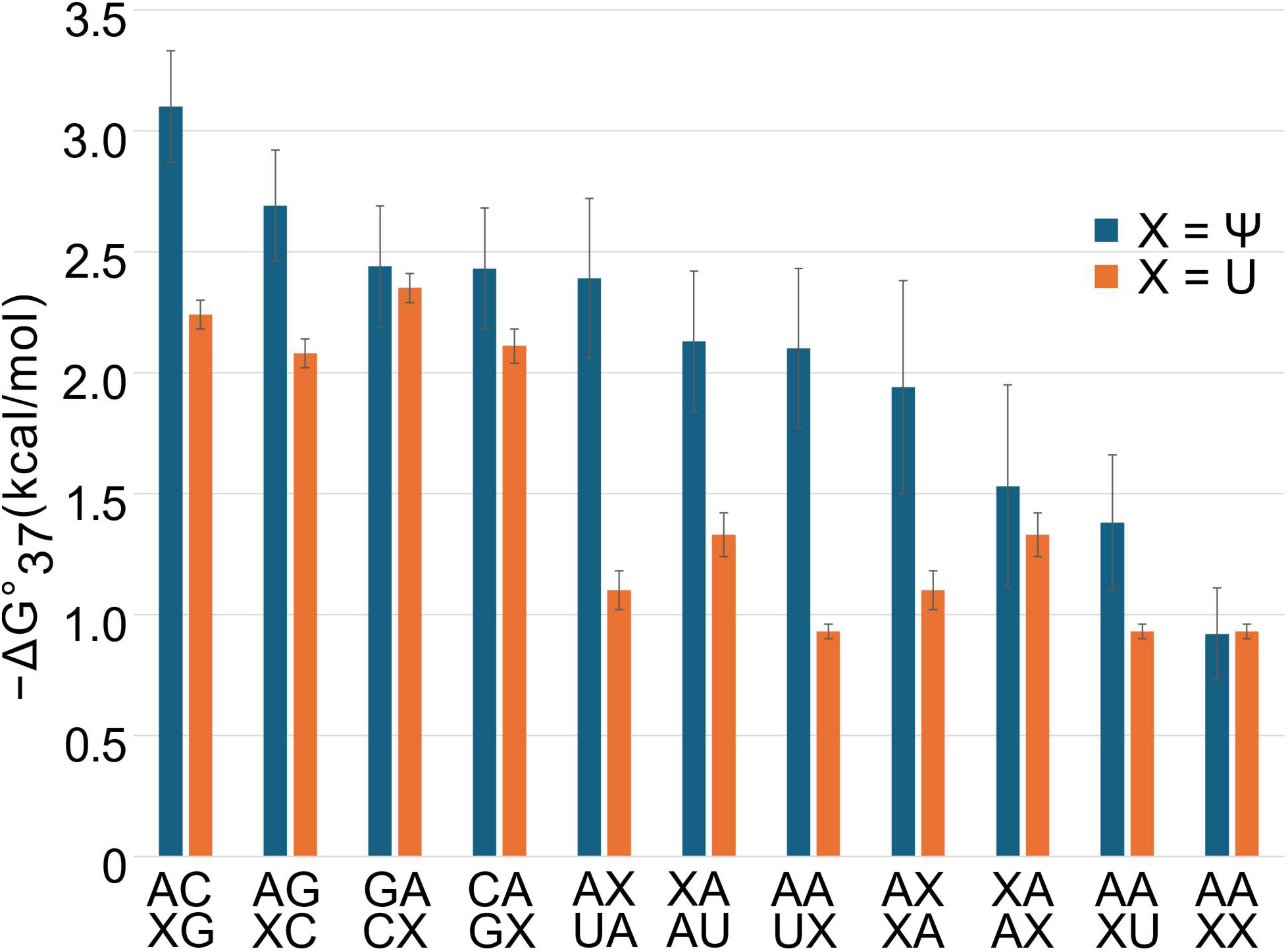
Ψ-A base pair stacking parameters are more stabilizing than analogous U-A base pair parameters. Stacking parameters were determined by linear regression, and the uncertainties are the standard errors of regression. The identity of X can either be Ψ (blue) or U (orange). On average, each Ψ-A substitution for a U-A is stabilizing by -0.56 ± 0.13 kcal/mol. This depends on sequence, and the stability estimates range from -3.1 ± 0.23 kcal/mol to -0.92 ± 0.19 kcal/mol.

Figure 3 compares Ψ-G stacks to the analogous U-G stacks. The stabilities for these stacks range from -3.21 ± 0.20 kcal/mol to 0.18 ± 0.29 kcal/mol. On average, each Ψ-G substitution for a U-G stack is stabilizing by -0.36 ± 0.11 kcal/mol. This stabilization is also sequence-context dependent, ranging from -3.0 ± 0.45 kcal/mol to 0.57 ± 0.30 kcal/mol (Supplementary Table S4). A subset of stacks that are substituted with Ψ are not very different from their U analogues (|ΔΔG°_37_|<0.25 kcal/mol), for example G-U followed by a G-Ψ pair and U-A followed by G-Ψ. There are two cases in which the base pair stacks are destabilizing (5′GU3′/3′UG5′ and 5′UG3′/3′GU5′), both with a free energy value of 0.72 ± 0.19 kcal/mol. Although it is important to note that the uncertainty in these measurements is large.

**Figure 3:**
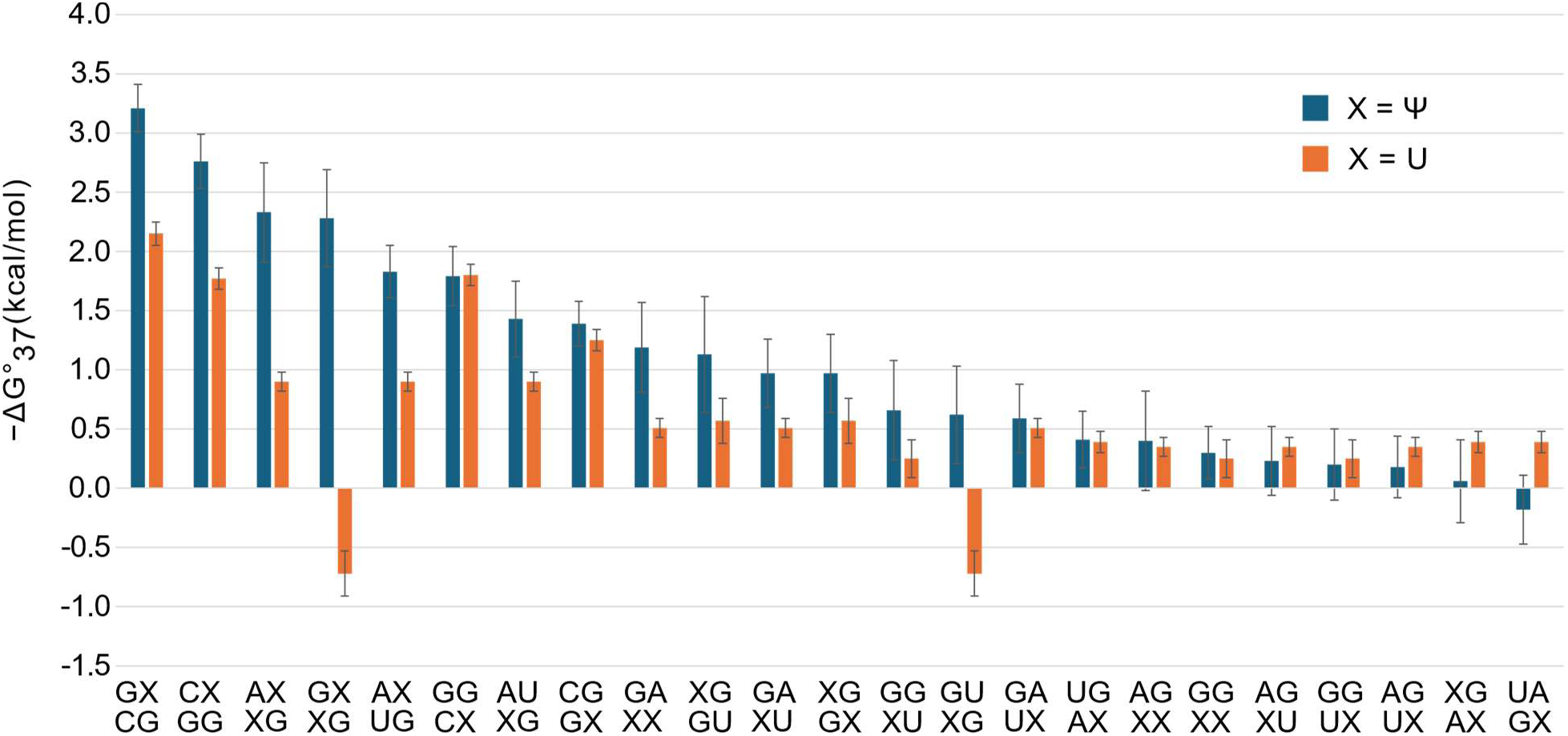
Ψ-G base pair stacking parameters are more stabilizing than analogous U-G base pair parameters. Stacking parameters were determined by linear regression, and the uncertainty errors are the standard errors of regression. On average, each Ψ-G substitution for a U-G is stabilizing by -0.36 ± 0.11 kcal/mol. This is sequence dependent, and the stability estimates range from -3.21 ± 0.20 kcal/mol to 0.18 ± 0.29 kcal/mol.

A terminal Ψ-A pair is about as destabilizing as a terminal U-A pair in the Turner 2004 model, with values 0.42 ± 0.21 kcal/mol and 0.45 ± 0.04 kcal/mol, respectively (Figure 4). Chen et al. [37] found that terminal U-G pairs are not destabilizing. However, we found that a terminal Ψ-G is destabilizing by 0.21 ± 0.15 kcal/mol. These comparisons are shown in Figure 4.

**Figure 4:**
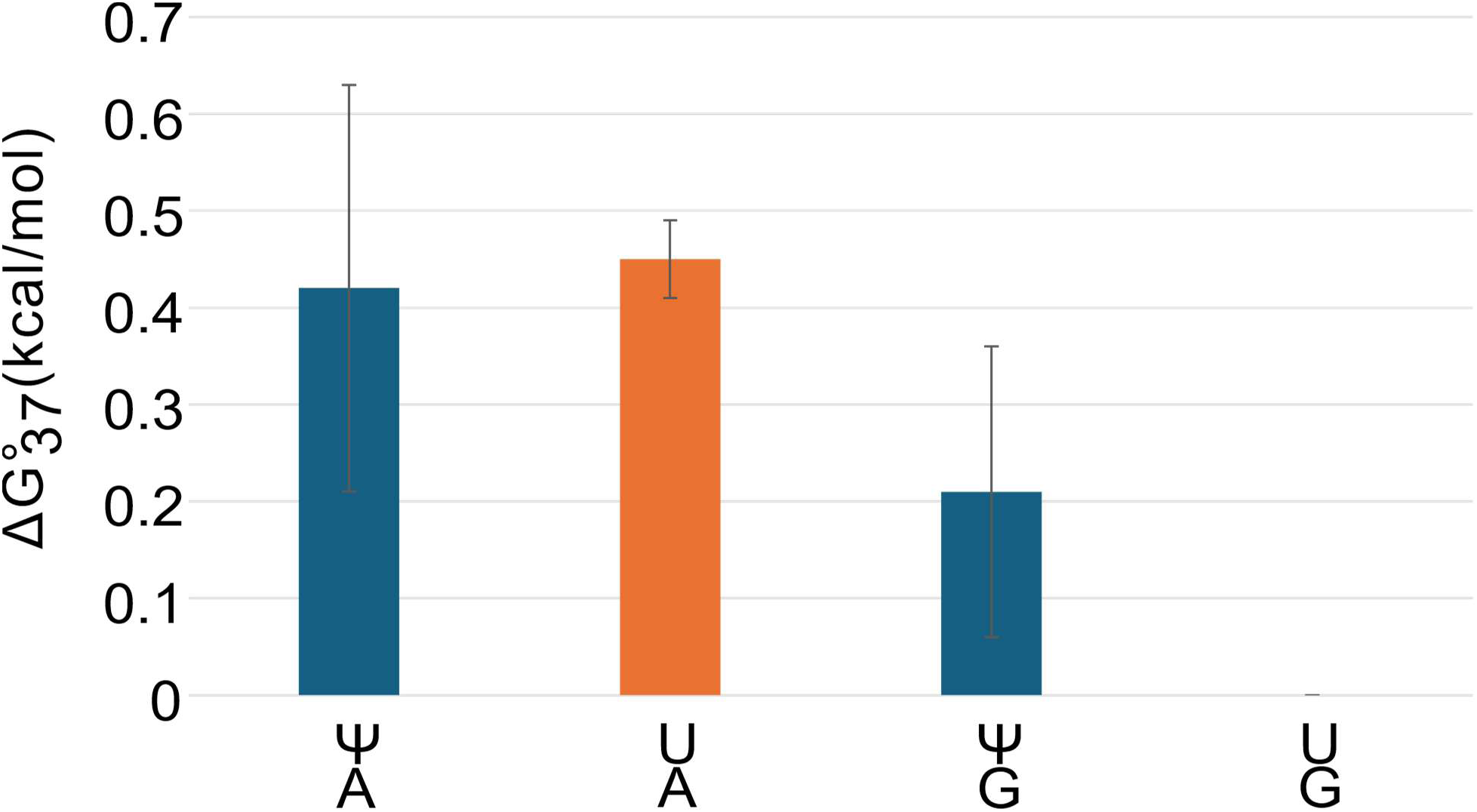
Terminal Ψ-A and Ψ-G pairs are destabilizing. These results are derived from linear regression, with uncertainty errors as the standard errors of regression. A Ψ-A at the end of a helix is comparable to having a U-A pair. Ψ-A terminal pairs are destabilizing by 0.42 ± 0.21 kcal/mol, while U-A terminal pairs are destabilizing by 0.45 ± 0.04 kcal/mol [48]. Terminal Ψ-G pairs are more destabilizing compared to U-G pairs, with a value of 0.21 ± 0.15 kcal/mol. The current model does not have a term for terminal U-G pairs because the stability cost is close to zero [37].

### Dangling ends and Terminal mismatches

Dangling ends, a single unpaired nucleotide adjacent to a helix, are known to stabilize RNA secondary structure. We studied both 5’ and 3’ dangling ends (optical melting experiment results in Supplementary Table S5). On average, 5’ and 3’ dangling ends with Ψ are more stabilizing compared to an analogous dangling end with U. This effect is sequence-context dependent (Supplementary Table S6). A 5’ Ψ dangling end is estimated and measured to be more stabilizing than an analogous U on G-C or U-A terminal pairs by an average of −0.44 ± 0.07 kcal/mol. A 3’ Ψ dangling end is also estimated and measured to be more stabilizing than an analogous U on C-G, G-C, and U-A by an average of −0.34 ± 0.06 kcal/mol.

Terminal mismatches are non-canonical pairs adjacent to helices that also stabilize secondary structure. We studied terminal mismatch duplexes containing Ψ within the mismatch pair and/or the adjacent base pair (optical melting results in Supplementary Table S7). Terminal mismatches are additionally stabilized when they contain a Ψ instead of an analogous U, and this effect is sequence-context dependent (Supplementary Table S8). For example, Ψ-Ψ mismatches next to A-U, G-C, and C-G terminal pairs are more stabilizing compared to the analogous U-U mismatches by -0.57 ± 0.20 kcal/mol. C-A and A-A mismatches next to A-Ψ are more stabilizing than the analogous A-U by -0.56 ± 0.23 kcal/mol.

### Bulge Loops

We performed 12 optical melting experiments to test the stability of bulge loops with Ψ (Supplementary Table S10). These sequences had a bulged Ψ, a Ψ located in a closing base pair, or a Ψ in the base pair adjacent to the closing pair. In the Turner 2004 parameters, bulge loop penalties are sequence independent, and the stability penalty grows with the size of the bulge loop [4]. The average ΔG°_37_ for a bulged Ψ (3.86 ± 0.26 kcal/mol) is close to the U value, (3.89 ± 0.62 kcal/mol). Because the values are consistent, we adopted the standard, sequence-independent value for a Ψ single bulge. Motif stabilities for bulge loops are in Supplementary Table S11.

### Internal loops

We studied 13 model internal loops by optical melting with Ψ in the loop or in a base pair closing the loop. Two additional internal loops had been previously studied [67] (Supplementary Table S9). The motif stabilities for internal loops are in Supplementary Table S12. Our analysis included, among others, Ψ-Ψ, Ψ-U, U-Ψ, and Ψ-C mismatches. The results show that these first mismatches stabilize internal loop formation compared to U-U first mismatches, where a first mismatch is defined as the non-canonical pair of nucleotides directly adjacent to the closing helix. Internal loops of 2×2 nucleotide or larger contain two first mismatches, one adjacent to each closing helix. Figure 5 shows example internal loops with Ψ-Ψ and Ψ-U mismatches.

**Figure 5:**
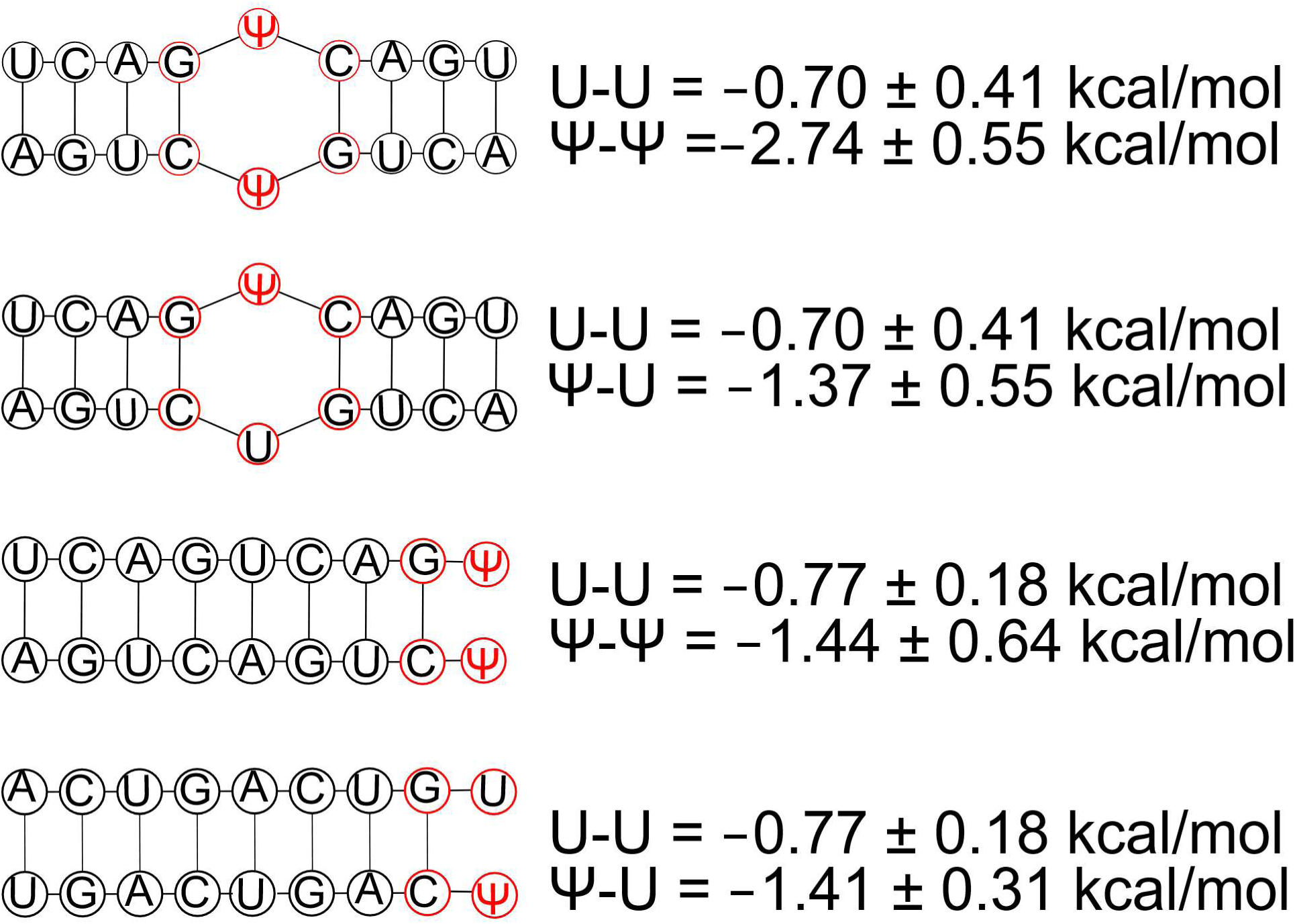
Pseudouridine is stabilizing as both internal and terminal mismatches. Uncertainty estimates are propagated from the uncertainty of the individual optical melting experiments as explained in the Materials and Methods. Pseudouridine is stabilizing as a Ψ-Ψ and Ψ-U single mismatch and as a Ψ-Ψ and U-Ψ terminal mismatch.

Based on an average of four experiments, the first Ψ-Ψ mismatch stabilizes by an additional -1.68 ± 0.35 kcal/mol. From five experiments, the first Ψ-U and U-Ψ mismatch stabilize by an additional -1.0 ± 0.34 kcal/mol. From a single experiment, the first Ψ-C mismatch stabilizes by an additional -0.68 ± 0.61kcal/mol. The average of two experiments shows that a Ψ-A or A-Ψ closing pair stabilizes by -0.47 ± 0.10 kcal/mol compared to an analogous U-A or A-U closing pair.

We observed that a 2×2 Ψ-Ψ mismatch is almost twice as stabilizing as a 1×1 Ψ-Ψ mismatch. This result reflects the stronger stacking stability of Ψ relative to U.

### Hairpin loops

We studied eight hairpin loops with Ψ in the loop or in the closing base pair by optical melting (Supplementary Table S13). In the Turner 2004 model, the stability of hairpin loops depends on the size of the loop and the sequence in the closing pair and first mismatch [36]. Here, we tested this model by placing Ψ in the closing pair, the first mismatch, or in the interior of the hairpin loop. The stability of hairpin loops containing Ψ are well modeled by the Turner 2004 model (Supplementary Table S14). In this model, there is a bonus of -0.95 ± 0.81 kcal/mol when a hairpin loop has a U-U first mismatch [4]. We applied this bonus to U-Ψ, Ψ-U, and Ψ-Ψ first mismatches because the model better predicted the experimental ΔG°_37_ when included.

For a subset of tetraloops (4-nucleotide hairpins), triloops (3-nucleotide hairpins), and hexaloops (6-nucleotide hairpins), the standard model does not work well, and a lookup table of experimental results is used for these [4]. For the Ψ alphabet in RNAstructure, hairpin loop tables for special triloops, tetraloops, and hexaloops remain largely unchanged. Three of the sequences that were measured in this study are special tetraloop sequences containing Ψ (Supplementary Table S13). The measured stability values for the sequences GGAC<u>ΨUCG</u>GUCC and GGAC<u>UΨCG</u>GUCC were not well described by the experimental value of the U analogue in the lookup table; the differences are -2.41 ± 0.50 kcal/mol and 1.21 ± 0.42 kcal/mol, respectively (Supplementary Table S14). The stability values for C<u>ΨUCG</u>G and C<u>UΨCG</u>G were therefore added to the tetraloop tables.

### Modeling alternative structures of *S. cerevisiae* U2 snRNA

Three pseudouridines, at positions 35, 42, and 44, are found in *S. cerevisiae* U2 snRNA under optimal growth conditions. There are two additional pseudouridines, at positions 56 and 93, that are induced by heat shock or nutrient deprivation [40]. It has been determined that U2 snRNA toggles between two alternative structures, stem IIa and stem IIc (Figure 6) [68,69]. Positions 56 and 93 lie within the region of stem II that can adopt alternative conformations, and these modifications affect the equilibrium between stem IIa and stem IIc [40]. Using single-molecule fluorescence resonance energy transfer (smFRET), the structure and dynamics of stem II in the presence and absence of the induced pseudouridines at positions 56 and 93 were investigated [70]. Pseudouridines at positions 56 and 93 stabilize the formation of the stem IIc structure.

**Figure 6:**
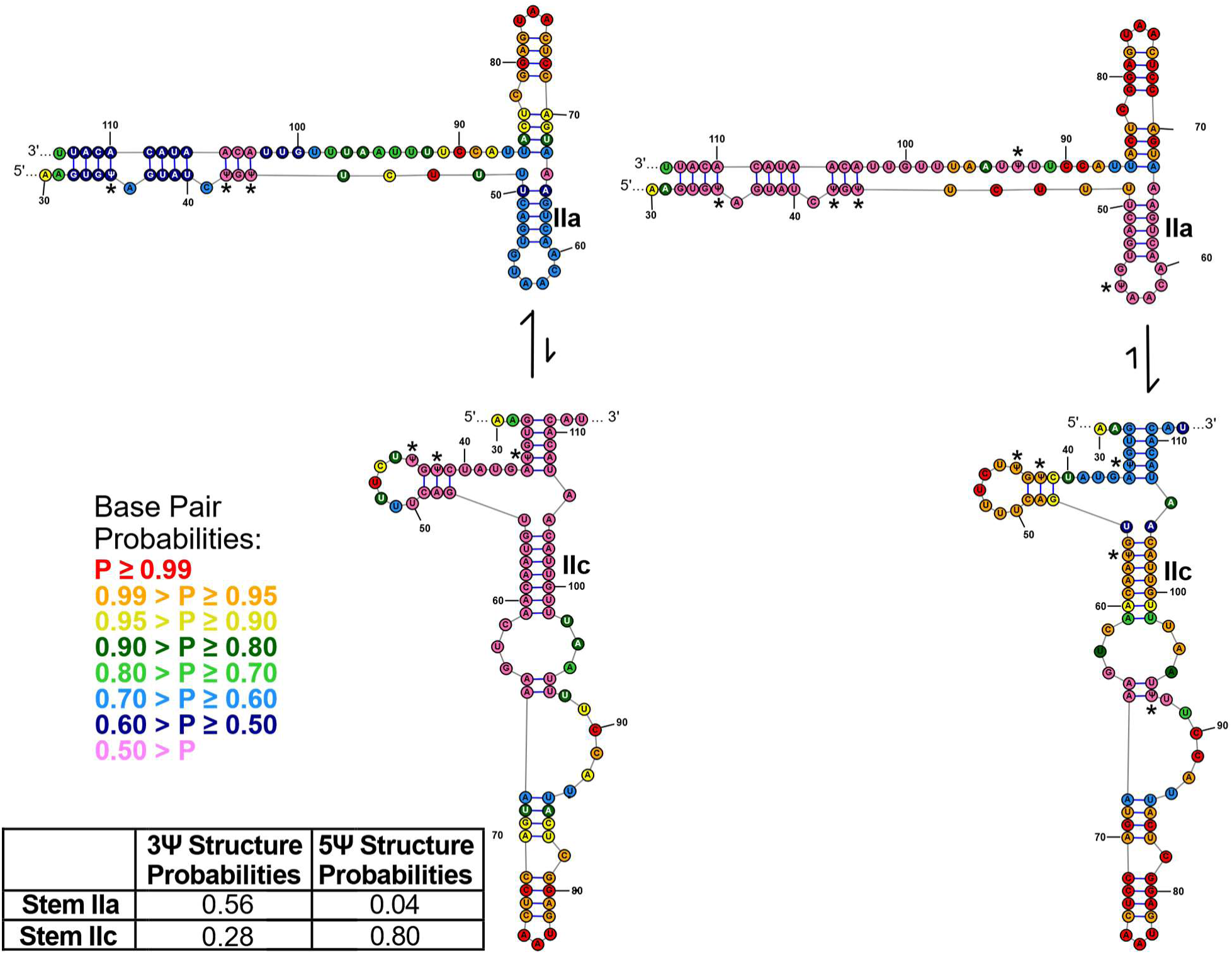
Alternative secondary structures of S. cerevisiae U2 snRNA predicted using pseudouridine parameters. The structure of U2 snRNA is compared between a sequence with three pseudouridines that are constitutively expressed (left) and a sequence with two additional pseudouridines that are induced by heat shock or nutrient deprivation (right). Positions of pseudouridines are marked with * to improve their visibility. The probability of the stem IIa forming is 0.56 in the three pseudouridine structure, and that decreases to 0.04 in the five pseudouridine structure. The probability of stem IIc formation increased from 0.28 in the three pseudouridine structure to 0.80 in the five pseudouridine structure.

We used RNAstructure to model the two alternative structures to test our parameters (Figure 6). We compared the structure with three pseudouridines to the structure with five pseudouridines. By predicting the maximum expected accuracy structure (RNAstructure *MaxExpect*) [71], we observe stem IIa when three pseudouridines are present, and stem IIc when the two inducible pseudouridines at positions 56 and 93 are added. We used RNAstructure’s *Probscan* [66] to quantify the probability of stem IIa and stem IIc forming in the two conformations. The probability of stem loop IIa is 0.56 in the three-pseudouridine structure but reduces to 0.04 in the five-pseudouridine structure. Conversely, the probability of stem IIc forming increases from 0.28 in the three-pseudouridine structure to 0.80 in the five-pseudouridine structure.

## Discussion

Given the widespread and important biological roles of Ψ, it is critical to understand the impact of pseudouridylation on RNA structure [20,21,24]. In this work, we derived folding free energy nearest neighbor parameters, a.k.a. Turner rules, for modeling the secondary structure and estimating the folding stability of RNA sequences containing the nucleotides A, C, G, U, and Ψ. The Turner rules for canonical base pairs were derived from 802 optical melting experiments [33]. Here, we extrapolate the parameters for Ψ using a set of 210 optical melting experiments. Prior work with m^6^A showed that a limited set of 45 optical melting experiments was adequate for extrapolating nearest neighbor terms [33,34]. Key to the selection of experimental sequences was a prior sensitivity analysis that identified the parameters with the greatest influence on secondary structure prediction precision [39]. This guided our focus on Ψ experiments for helical stacking parameters, dangling ends, terminal mismatches, internal loops, hairpin loops and bulges.

### Ψ stabilizes RNA folding

Hudson et al. (2017) previously reported eight helical stacking parameters for Ψ-A pairs based on stacks containing a single Ψ-A substitution for a U-A pair [38]. These parameters and the new parameters reported here, on average, agree within 0.30 kcal/mol RMSD of each other (with largest percent difference of 26 %) (Figure 7A and Supplementary Table S14). Mauger et al (2019) reported fully substituted stacks in which all U nucleotides were replaced by Ψ [10]; those parameters agree with ours within 0.38 kcal/mol RMSD (largest percent difference was 28 %) (Figure 7B and Supplementary Table S15). This work incorporates single substituted stacks, fully substituted stacks, stacks containing Ψ-G pairs, and loop parameters. The new parameters, however, have larger standard errors despite being fit with more data, likely reflecting the added variability introduced by simultaneously fitting both Ψ-G and Ψ-A stacks.

**Figure 7:**
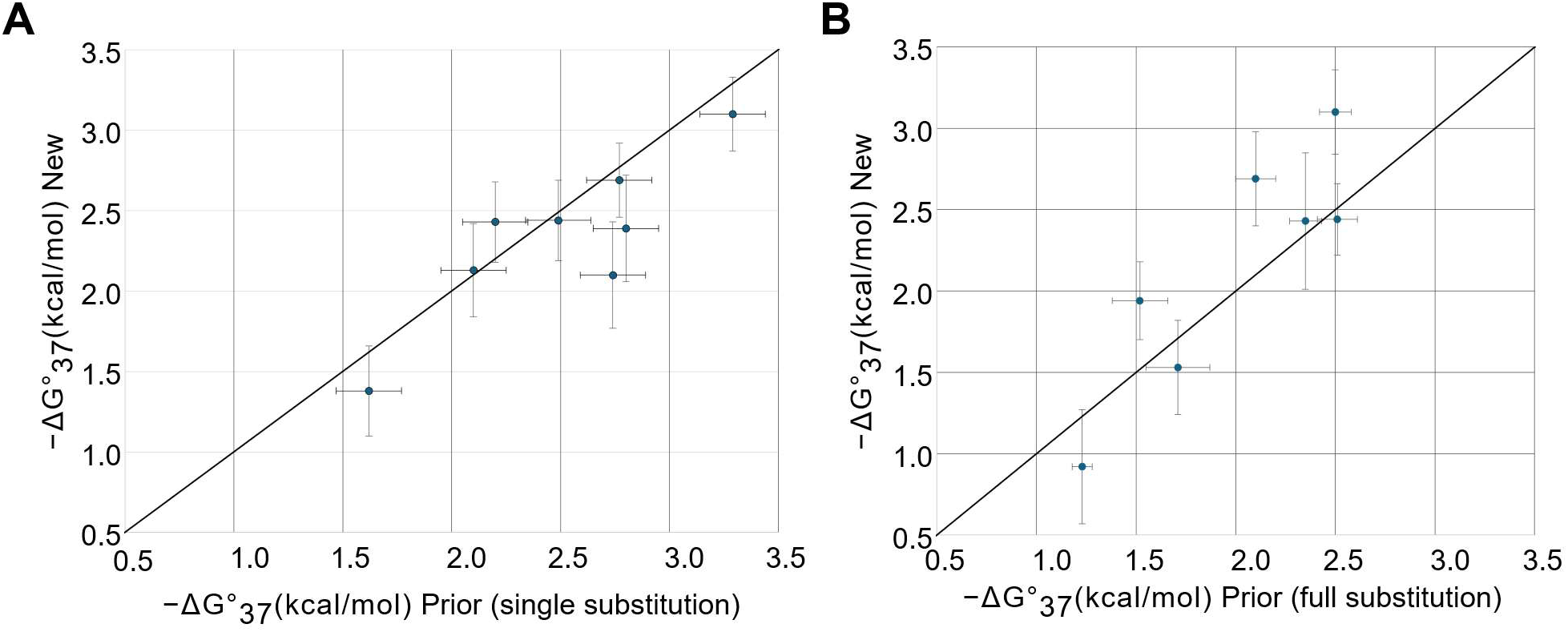
The new Ψ–A stacking parameters are comparable to those reported previously. Panel (**A**) compares the new parameters with those reported for single-substitution stacks [38]. The RMSD for ΔΔG°37 between the two datasets is 0.30 kcal/mol, and the diagonal line is shown, which would indicate perfect agreement. Panel (**B**) compares the new parameters with those reported for fully substituted stacks, in which all U nucleotides were replaced by Ψ [10]. The RMSD for ΔΔG°37 in this comparison is 0.38 kcal/mol, and the diagonal line is shown for reference.

Overall, substituting U with Ψ enhances folding stability in both helical stacks and loop regions, although the magnitude of this effect is sequence-context dependent. For internal loops, the Ψ-Ψ first mismatch pairs were found to be more stabilizing than U-U, with a stabilizing energy of -1.68±0.35 kcal/mol, while Ψ-U and U-Ψ pairs showed stabilization of -1.00±0.34 kcal/mol. A Ψ-C first mismatch for 1×1nucleotide internal loops was stabilizing by -0.68±0.61 kcal/mol. An A-Ψ closing pair provides additional stability of -0.48±0.10 kcal/mol. Terminal mismatches with Ψ were also stabilizing relative to their U analogues. Figure 5 illustrates example Ψ-Ψ and Ψ-U terminal mismatches that were analyzed. Both 3’ and 5’ dangling ends with Ψ were stabilizing relative to their analogous U counterparts. A single Ψ bulge showed comparable stability to a single U bulge, although one outlier was observed. Hairpin loops generally conform to the Turner 2004 model, with the exception of specific tetraloop sequences, C<u>ΨUCG</u>G and C<u>UΨCG</u>G.

In the Turner 2004 parameters, U-A base pairs incur a stability penalty of 0.45 ± 0.04 kcal/mol when they appear at the end of a helix (called terminal pairs)[48]. Terminal Ψ-A pairs were found to be almost as destabilizing as U-A pairs at 0.42 ± 0.21 kcal/mol. Interestingly, while terminal G-U pairs have no penalty [37], it costs free energy to have a Ψ-G terminal pair at 0.21 ± 0.15 kcal/mol. These differences suggest that the terminal pairs are behaving differently when <u>Ψ is</u> <u>substituted for U. More recent helical terms than the 2004 Turner model consider the terminal and</u> <u>penultimate pairs when determining terminal base pair terms</u> [72]<u>. Future work with additional</u> <u>experiments could use the last two terminal pairs to determine end terms.</u>

The stabilizing effect of Ψ is primarily associated with differences in enthalpy. For 51 analogous helix sequences (marked in Supplementary Table S2 and a sequence from [58]), we plotted the enthalpy and entropy values of Ψ-containing helices against those containing uridine (Figure 8). Paired t-tests showed statistically significant differences for both parameters (p = 0.0003 for ΔH° and p = 0.007 for ΔS). Duplexes containing pseudouridine exhibit lower ΔH° and lower ΔS° values. A lower ΔH° is stabilizing, whereas a lower ΔS° is destabilizing. Consistent with these trends, the ΔG°_37_ values (Supplementary Figure S2) follow the same pattern, with a significant paired t-test p-value of 3.4 × 10⁻¹⁵, further indicating that the increased stability of Ψ-containing sequences is driven by the enthalpic contribution.

**Figure 8.**
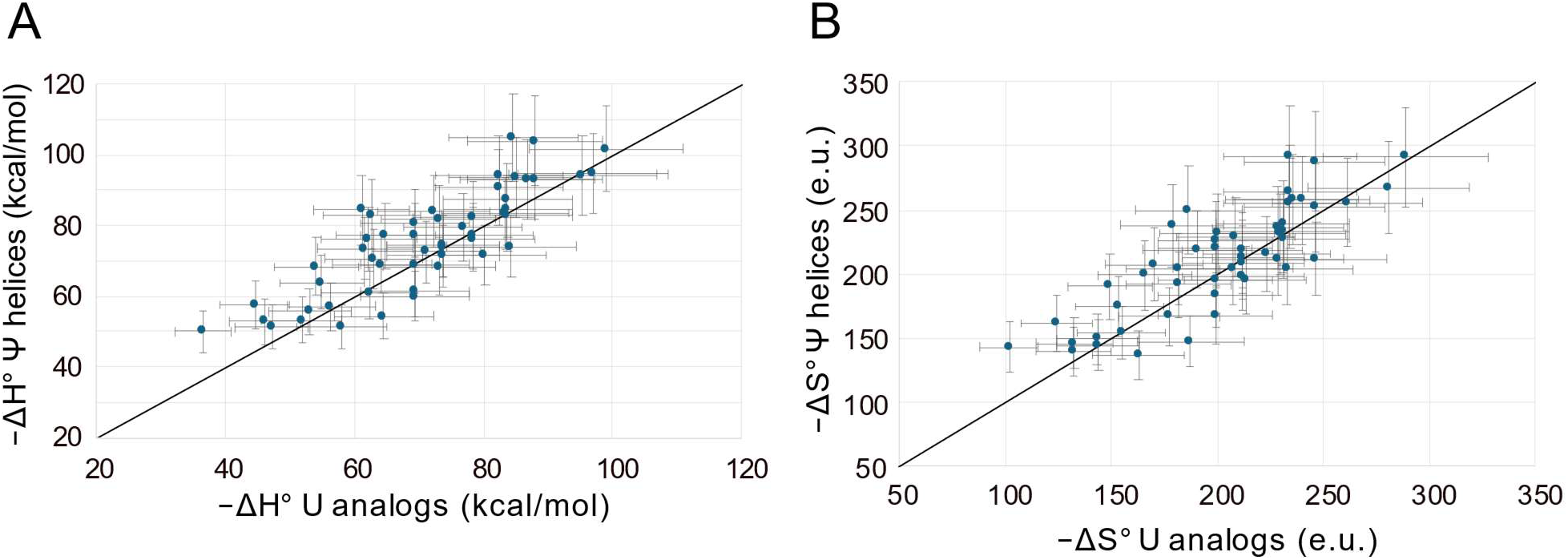
ΔH° and ΔS° are lower for Ψ-containing helices compared to their analogous U-containing duplexes. Panel (**A**) shows the enthalpy (ΔH°) values for the same 51 analogous duplexes plotted for Ψ-containing duplexes versus U-containing analogs, with a significant paired t-test difference (p = 0.0003). The diagonal line is shown for reference. Panel (**B**) shows the entropy (ΔS°) values for 51 analogous duplex sequences plotted for Ψ-containing helices versus their U-containing analogues. Paired t-tests showed a statistically significant difference (p = 0.007). The diagonal line is shown for reference. In both panels, Ψ-containing duplexes exhibit lower ΔH° and lower ΔS° values relative to their U-containing analogs. Experimental errors for ΔH° and ΔS° were estimated as 12% and 13.5%, respectively, as described in [48].

The molecular origin of the enhanced enthalpy is not clear at this time but can come from any source of new molecular interactions including stacking and hydrogen bonding, with the latter being afforded by the donation ability of the N1H, now opposite of the WCF face. The N1H has myriad potential acceptors, which outnumber donors 5:1 in RNA, including the bases, sugars and phosphates [73].

### Applications

These parameters improve the modeling of RNA structures containing Ψ and enable the estimation of the stability of structures directly from the primary sequence. We demonstrated the utility by modeling the alternative secondary structures of U2 snRNA. We used the parameters to quantify the probability of forming either stem IIa or stem IIc when the structure contains three pseudouridines or five pseudouridines. These structures were previously studied using smFRET and RNase T1 structure probing [70]. The parameters allow us to estimate the probabilities of the alternative structures that form, providing a quantitative framework. This serves as one example of how the parameters can be applied to develop structural hypotheses.

### Parameter availability

The RNAstructure software package was updated to incorporate Ψ nearest neighbor parameters, starting with version 6.6. These parameters are available as plain text files and can be accessed at https://rna.urmc.rochester.edu/RNAstructure.html. Parameters sets are provided for both fully substituted sequences, where Ψ replaces each U, and sequences containing a mixture of U and Ψ. For fully substituted sequences, calculations are faster when using the fully substituted parameters because the set of parameters is smaller.

## Data Availability

The data are available in the article and the supplementary materials.

## Acknowledgements

We thank Yi-Tao Yu for helpful conversations about the U2 snRNA.

## Author Contributions

Thandolwethu S. Shabangu (formal analysis, investigation, writing -original draft), Elzbieta Kierzek (conceptualization, funding acquisition, investigation, writing -review & editing, supervision), Sebastian Arteaga (investigation), Gregory S. Orf (investigation) Julia Stone (Investigation), Olivia M. Hiltke (formal analysis, writing -review & editing), Megan Miaro (supervision, formal analysis, writing -review & editing), Elizabeth A. Jolley (investigation, writing -review & editing), Marta Soszyńska-Jóźwiak (investigation), Marta Szabat (investigation), Sharon Aviran (conceptualization, formal analysis), Philip C. Bevilacqua (supervision, investigation, writing -review & editing), Brent M. Znosko (conceptualization, supervision, funding acquisition, writing -review & editing), Ryszard Kierzek (conceptualization, funding acquisition, investigation, writing -review & editing, supervision), David H. Mathews (conceptualization, funding acquisition, writing -review & editing, formal analysis, supervision, software)

