## Supplemental Figures and Tables for "Nearest Neighbor Parameters for Estimating the Folding Stability of RNA Including Pseudouridine"

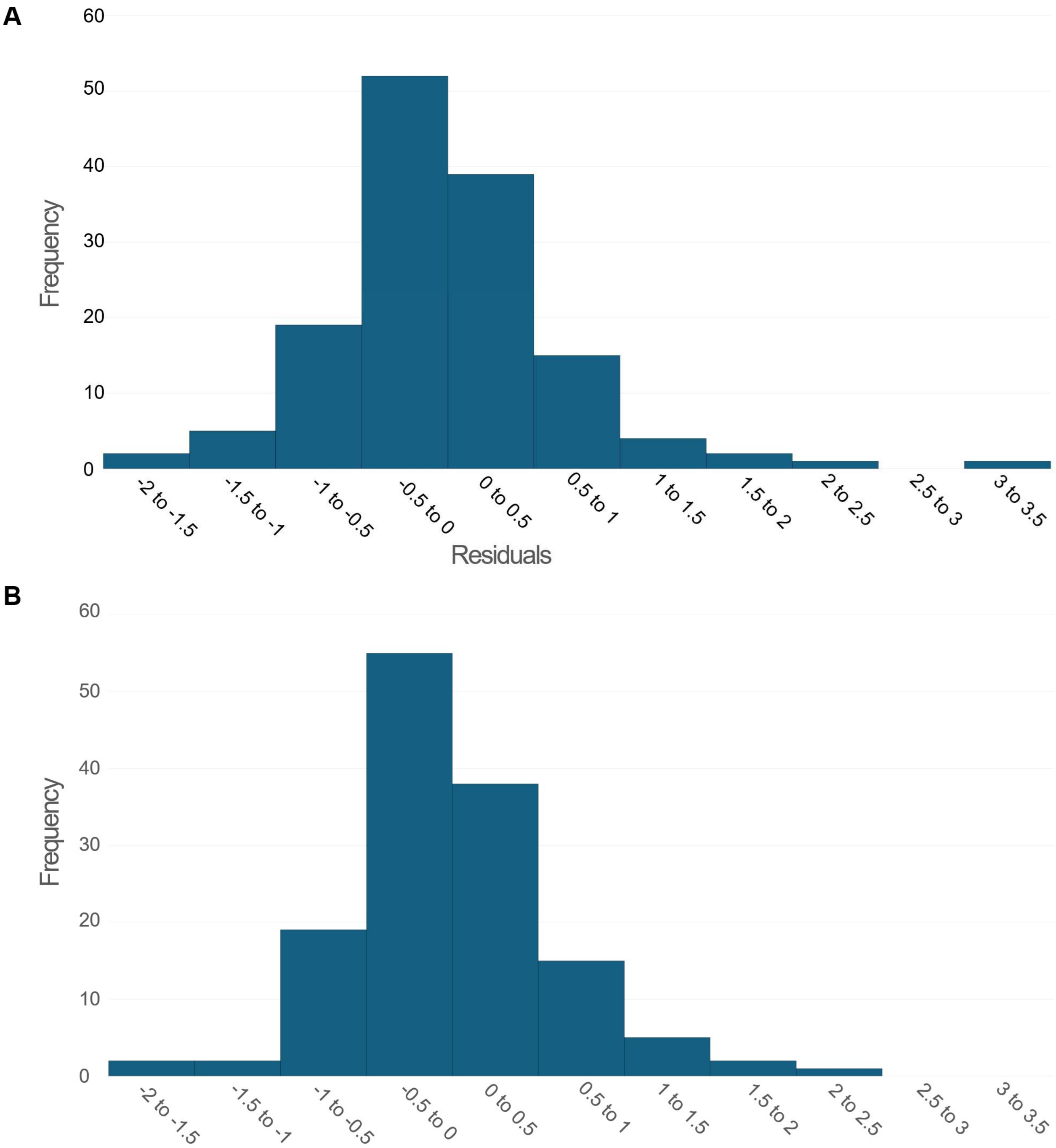

**Figure S1:** Histograms of residuals from the linear regression fit. Plot A shows the distribution of residuals for all duplexes that fit the two-state model. One duplex, 5’GCUΨCGC/3’CGAGGCG, was identified as an outlier and excluded from the analysis. Plot B shows the residual distribution after the outlier sequence was removed and the data was refit.

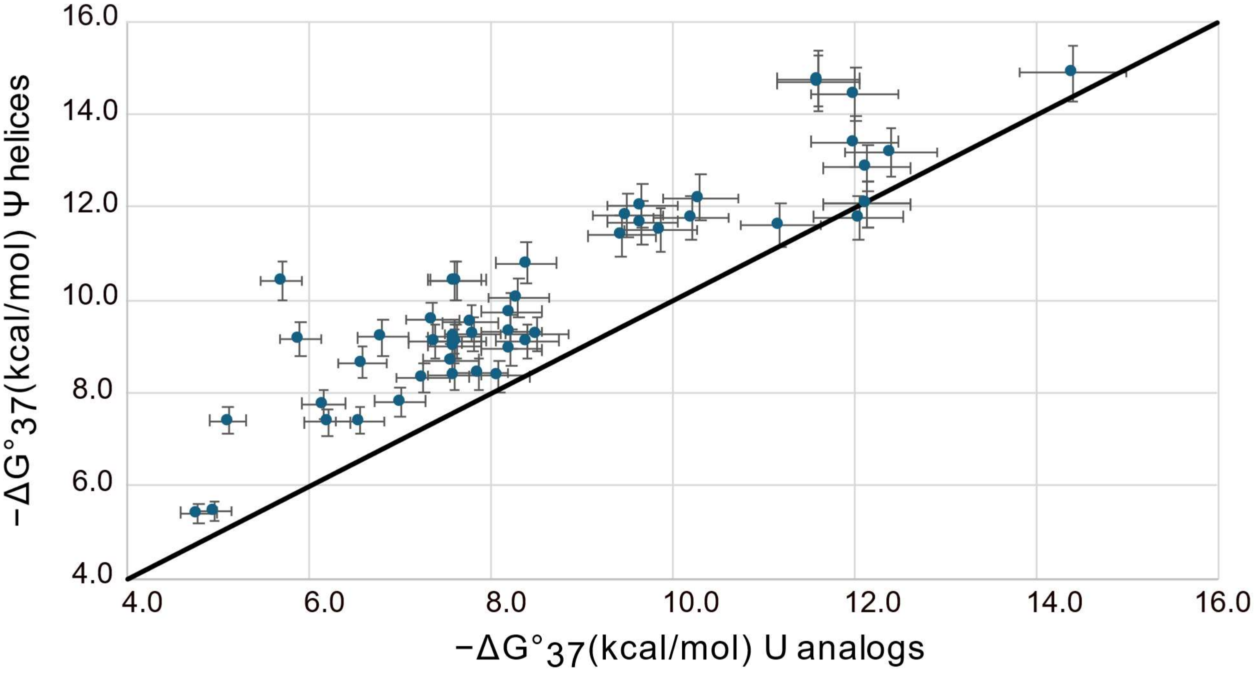

**Figure S2:** $\Delta G°_{37}$ values are lower for Ψ-containing helices compared to their analogous U-containing duplexes. The $\Delta G°_{37}$ values for the 51 analogous duplex sequences show a significant paired t-test difference ($p = 3.4 \times 10^{-15}$). Experimental errors were estimated as 4%, as described in [48] .

**Table S1**. Agreement between calibration duplex optical melting data. The buffer used was 1 M sodium chloride, 20 mM sodium cacodylate, 0.5 mM $Na_2EDTA$, pH 7.

| Site: | Duplex | Average of curve fits | | | | | $T_M^{-1}$ vs log $C_T$ plots | | | |
|---|---|---|---|---|---|---|---|---|---|---|
| | | -ΔH° (kcal/mol) | -ΔS° (eu) | -ΔG°$_{37}$ (kcal/mol) | $T_M$[a] (˚C) | | -ΔH° (kcal/mol) | -ΔS° (eu) | -ΔG°$_{37}$ (kcal/mol) | $T_M$[a] (˚C) |
| Poznan[51] | CGCGCG GCGCGC | 58.4±5.3 | 157.5±16.2 | 9.58±0.33 | 59.0 | | 56.0±2.8 | 150.3±10.8 | 9.42±0.16 | 59.0 |
| St. Louis [51] | CGCGCG GCGCGC | 60.2±3.6 | 164.2±11.2 | 9.29±0.15 | 56.7 | | 62.9±3.0 | 172.6±9.1 | 9.39±0.16 | 56.3 |
| Poznan (2022)[33] | CGCGCG GCGCGC | | | | | | 47.8±1.9 | 124.0±5.9 | 9.38±0.14 | 62.8 |
| Rochester (1985) [55] | CGCGCG GCGCGC | | | | | | 54.5 | 146.4 | 9.12 | 57.8 |
| Poznan[51] | GCUACG CGAUGC | 57.8±3.0 | 161.5±9.5 | 7.74±0.17 | 43.5 | | 56.4±3.7 | 157.2±11.5 | 7.68±0.11 | 43.2 |
| St. Louis[51] | GCUACG CGAUGC | 63.1±8.8 | 179.4±27.9 | 7.69±0.12 | 41.6 | | 58.9±5.3 | 166.3±17.0 | 7.36±0.13 | 41.2 |
| Rochester (1998) [48] | GCUACG CGAUGC | | | | | | 58.0 | 162.7 | 7.56 | 42.5 |
| Poznan[51] | GCGUUCGC CGCUUGCG | 68.8±6.3 | 196.3±19.4 | 7.98±0.27 | 47.4 | | 65.5±3.0 | 186.0±9.3 | 7.84±0.08 | 47.5 |
| St. Louis[51] | GCGUUCGC CGCUUGCG | 75.6±5.1 | 218.8±15.5 | 7.69±0.12 | 45.7 | | 73.9±3.4 | 213.7±10.8 | 7.62±0.07 | 45.4 |
| Rochester (1995) [54] | GCGUUCGC CGCUUGCG | 70.1±3.8 | 200.7±11.9 | 7.88±0.17 | 46.9 | | 65.5±0.9 | 186.5±2.9 | 7.69±0.03 | 46.7 |

[a] Tm calculated for $10^{-4}$ M oligomer concentration.

**Table S2.** Optical melting experiment data for helices used to fit nearest neighbor parameters for Ψ-A and Ψ-G base pair stacks. ΔH° is the folding enthalpy change, ΔS° is the folding entropy change, and $\Delta G°_{37}$ is the folding free energy change at 37 °C, determined from the enthalpy and entropy changes. $T_M$ is the melting temperature. An additional 24 duplexes, as reported by [38], and one from [58], were incorporated to fit the data . The pseudouridine base pairs are shown in red and their nearest neighbors are shown in blue.

| Duplexes (5'-3') | Average of curve fits | | | | $T_M^{-1}$ vs log $C_T$ plots | | | |
|---|---|---|---|---|---|---|---|---|
| | -ΔH° (kcal/mol) | -ΔS° (eu) | $-\Delta G°_{37}$ (kcal/mol) | $T_M^b$ (°C) | -ΔH° (kcal/mol) | -ΔS° (eu) | $-\Delta G°_{37}$ (kcal/mol) | $T_M^b$ (°C) |
| CCΨAGG[c,f]<br>GGAΨCC | 59.7±4.0 | 163.8±12.5 | 8.89±0.15 | 54.6 | 68.3±3.4 | 190.5±10.3 | 9.26±0.16 | 54.2 |
| GCΨACG[c,f]<br>CGAΨGC | 55.4±3.1 | 149.9±9.2 | 8.87±0.25 | 50.7 | 51.2±4.2 | 137.1±13.2 | 8.71±0.17 | 50.8 |
| GCAΨCG[c,f]<br>CGΨAGC | 57.2±7.4 | 157.6±23.0 | 8.38±0.33 | 47.4 | 53.2±4.1 | 144.5±12.3 | 8.33±0.13 | 47.8 |
| GGCGCΨ[c,f]<br>ΨCGCGG | 58.7±3.5 | 159.6±10.6 | 9.23±0.23 | 57.0 | 57.1±4.4 | 154.7±13.4 | 9.13±0.21 | 57.0 |
| GCGΨGC[c,f]<br>CGΨGCG | 51.9±5.6 | 143.3±17.5 | 7.46±0.14 | 48.0 | 52.8±1.1 | 146.3±3.4 | 7.41±0.02 | 47.5 |
| GCAΨGC[c,f]<br>CGΨACG | 64.2±4.0 | 177.0±11.7 | 9.29±0.33 | 55.5 | 60.9±5.1 | 167.1±15.6 | 9.09±0.26 | 55.4 |
| GGΨACC[c,f]<br>CCAΨGG | 62.7±4.2 | 171.4±12.8 | 9.56±0.22 | 57.5 | 63.8±4.4 | 174.9±13.4 | 9.58±0.24 | 57.2 |
| GCGCAΨ[g]<br>CGCGUG | 69.8 ± 0.5 | 190.7 ± 1.6 | 10.66 ± 0.02 | 56.5 | 69.2 ± 0.7 | 189.0 ± 2.2 | 10.62 ± 0.04 | 56.5 |
| GCGCCΨ[g]<br>CGCGGG | 77.1 ± 0.5 | 213.9 ± 1.5 | 10.77 ± 0.02 | 55.1 | 76.9 ± 0.9 | 213.1 ± 2.7 | 10.76 ± 0.05 | 55.1 |
| GCGCGΨ[g]<br>CGCGCG | 86.8 ± 0.8 | 240.7 ± 2.5 | 12.13 ± 0.01 | 58.4 | 87.0 ± 1.9 | 241.3 ± 5.6 | 12.14 ± 0.12 | 58.4 |
| GCGCUΨ[g]<br>CGCGAG | 63.6 ± 0.2 | 177.6 ± 0.8 | 8.56 ± 0.01 | 47.2 | 63.6 ± 0.3 | 177.6 ± 0.8 | 8.56 ± 0.01 | 47.2 |
| ΨAGCGC[g]<br>GUCGCG | 57.2 ± 3.1 | 158.2 ± 9.9 | 8.12 ± 0.04 | 45.9 | 57.3 ± 1.5 | 158.5 ± 4.7 | 8.15 ± 0.04 | 46.0 |
| ΨCGCGC[g]<br>GGCGCG | 51.3 ± 5.3 | 141.0 ± 16.1 | 7.62 ± 0.03 | 43.8 | 55.6 ± 4.0 | 154.5 ± 12.7 | 7.67 ± 0.12 | 43.5 |
| ΨGGCGC[g]<br>GCCGCG | 52.2 ± 2.0 | 143.6 ± 6.5 | 7.65 ± 0.02 | 43.8 | 53.6 ± 0.7 | 148.3 ± 2.1 | 7.66 ± 0.01 | 43.7 |
| ΨUGCGC[g]<br>GACGCG | 56.4 ± 0.9 | 163.6 ± 3.1 | 5.67 ± 0.02 | 32.3 | 57.0 ± 1.1 | 165.4 ± 3.6 | 5.66 ± 0.02 | 32.4 |
| ΨGGCCG[c,f]<br>GCCGGΨ | 52.1±1.8 | 138.6±5.6 | 9.09±0.07 | 58.7 | 55.6±2.4 | 149.4±7.3 | 9.27±0.12 | 58.4 |
| ΨUGCAG[e,f]<br>GACGUΨ | 37.2±0.6 | 104.8±2.4 | 4.73±0.18 | 29.3 | 44.1±5.4 | 127.8±18.0 | 4.47±0.24 | 28.8 |
| UCΨAGA[c,f]<br>AGAΨCU | 46.3±3.3 | 131.5±10.9 | 5.49±0.18 | 35.7 | 49.9±6.5 | 143.2±21.3 | 5.46±0.24 | 35.6 |

| | | | | | | | | |
|---|---|---|---|---|---|---|---|---|
| CΨGCΨGG[e,f]<br>GAUGAUC | 54.3±4.4 | 149.8±14.0 | 7.84±0.53 | 44.7 | 39.2±6.4 | 101.6±20.2 | 7.65±0.43 | 46.1 |
| CΨAΨGAΨG[f]<br>GAΨAUΨAC | 58.6±2.6 | 165.8±8.1 | 7.13±0.18 | 40.2 | 60.9±5.0 | 173.4±16.2 | 7.11±0.12 | 40.0 |
| CGΨΨACG[f]<br>GCAAUGC | 63.3±2.6 | 174.3±7.9 | 9.23±0.16 | 50.8 | 60.2±1.3 | 164.7±4.1 | 9.11±0.04 | 50.9 |
| CGAΨACG[f]<br>GCΨAΨGC | 64.0±7.3 | 173.8±22.2 | 10.09±0.38 | 55.2 | 63.9±1.4 | 173.8±4.2 | 9.99±0.06 | 54.8 |
| CUCGCUC[c,f]<br>GAGΨGAG | 68.3±3.8 | 189.5±11.9 | 9.53±0.21 | 51.2 | 69.0±4.8 | 191.7±14.9 | 9.51±0.18 | 51.0 |
| GCAΨAGC[g]<br>CGUGUCG | 80.0 ± 5.6 | 228.2 ± 17.4 | 9.25 ± 0.24 | 47.9 | 79.8 ± 5.2 | 227.6 ± 16.3 | 9.23 ± 0.19 | 47.8 |
| GCAΨCGC[g]<br>CGUGGCG | 82. 3 ± 7.0 | 228.0 ± 21.1 | 11.55 ± 0.47 | 57.5 | 82.6 ± 3.6 | 229.2 ± 11.1 | 11.56 ± 0.21 | 57.1 |
| GCAΨGGC[g]<br>CGUGCCG | 82.5 ± 13.5 | 228.2 ± 41.3 | 11.72 ± 0.77 | 57.8 | 85.3 ± 8.3 | 237.2 ± 25.2 | 11.77 ± 0.51 | 57.3 |
| GCAΨUGC[g]<br>CGUGACG | 66.5 ± 2.2 | 186.0 ± 7.1 | 8.87 ± 0.11 | 48.3 | 74.9 ± 5.0 | 212.3 ± 15.7 | 9.08 ± 0.15 | 47.9 |
| GCCΨAGC[g]<br>CGGGUCG | 75.6 ± 8.0 | 206.9 ± 24.6 | 11.41 ± 0.38 | 58.4 | 76.0 ± 7.0 | 208.2 ± 21.4 | 11.44 ± 0.40 | 58.4 |
| GCCΨCGC[g]<br>CGGGGCG | 74.2 ± 6.9 | 196.7 ± 20.3 | 13.24 ± 0.60 | 67.8 | 78.0 ± 9.8 | 207.8 ± 29.0 | 13.51 ± 0.83 | 67.5 |
| GCCΨGGC[g]<br>CGGGCCG | 71.0 ± 3.7 | 186.7 ± 10.9 | 13.12 ± 0.37 | 68.7 | 78.5 ± 6.7 | 208.8 ± 19.9 | 13.73 ± 0.58 | 68.3 |
| GCCΨUGC[g]<br>CGGGACG | 73.7 ± 4.3 | 200.6 ± 13.0 | 11.42 ± 0.27 | 59.0 | 82.5 ± 5.0 | 227.5 ± 15.3 | 11.92 ± 0.30 | 58.7 |
| GCGΨAGC[g]<br>CGCGUCG | 75.5 ± 12.6 | 207.1 ± 37.9 | 11.27 ± 0.84 | 57.8 | 74.6 ± 9.9 | 204.2 ± 30.1 | 11.25 ± 0.69 | 58.0 |
| GCGΨCGC[g]<br>CGCGGCG | 71.7 ± 1.6 | 190.3 ± 5.1 | 12.71 ± 0.17 | 66.2 | 69.0 ± 9.1 | 182.2 ± 27.2 | 12.51 ± 0.78 | 66.4 |
| GCGΨGGC[g]<br>CGCGCCG | 73.3 ± 9.1 | 195.4 ± 27.3 | 12.73 ± 0.61 | 65.6 | 72.6 ± 7.0 | 193.1 ± 20.9 | 12.72 ± 0.57 | 65.9 |
| GCGΨUGC[g]<br>CGCGACG | 66.9 ± 5.7 | 181.9 ± 17.4 | 10.51 ± 0.29 | 56.6 | 69.4 ± 3.2 | 189.6 ± 9.9 | 10.62 ± 0.17 | 56.4 |
| GCUΨAGC[g]<br>CGAGUCG | 52.8 ± 18.2 | 145.9 ± 57.2 | 7.56 ± 0.49 | 43.1 | 48.2 ± 9.6 | 131.0 ± 30.3 | 7.61 ± 0.62 | 44.1 |
| GCUΨCGC[a,g]<br>CGAGGCG | 54.6 ± 8.9 | 157.1 ± 29.2 | 5.92 ± 0.26 | 33.6 | 57.2 ± 5.6 | 165.6 ± 18.3 | 5.78 ± 0.21 | 33.0 |
| GCUΨGGC[g]<br>CGAGCCG | 72.7 ± 4.2 | 198.6 ± 12.8 | 11.09 ± 0.29 | 57.7 | 75.5 ± 7.0 | 207.3 ± 21.3 | 11.22 ± 0.41 | 57.5 |
| GCUΨUGC[g]<br>CGAGACG | 67.2 ± 8.3 | 188.8± 26.3 | 8.60 ± 0.18 | 46.9 | 65.5 ± 4.4 | 183.8 ± 13.9 | 8.54 ± 0.12 | 46.8 |
| ΨΨGCAGG[g]<br>GGCGUCC | 53.4 ± 4.7 | 144.1 ± 15.1 | 8.66 ± 0.12 | 49.9 | 56.2±2.1 | 153.3±6.7 | 8.68±0.06 | 49.3 |

| | | | | | | | | |
|---|---|---|---|---|---|---|---|---|
| ΨΨGGAGG[g]<br>AACCUCC | 57.2 ±8.8 | 153.9 ± 26.6 | 9.43 ±0.68 | 53.6 | 58.8±9.5 | 159.2±29.2 | 9.42±0.68 | 53.1 |
| ACUΨAAGU[g]<br>UGAAΨUCA | 62.4 ±5.4 | 177.4 ± 17.2 | 7.38 ± 0.07 | 45.7 | 62.2±1.5 | 177.0±4.9 | 7.31± 0.02 | 45.4 |
| ACΨΨAAGU[c,f]<br>UGAAΨΨCA | 58.5±7.6 | 162.7±23.5 | 8.08±0.30 | 50.3 | 51.1±1.2 | 139.7±3.7 | 7.77±0.03 | 50.2 |
| AGAΨAΨCU[c,f]<br>UCΨAΨAGA | 59.3±5.7 | 162.4±17.4 | 8.92±0.29 | 54.9 | 54.4±1.3 | 147.6±3.8 | 8.65±0.05 | 54.9 |
| CCUGΨAGG[c,f]<br>GGAΨGUCC | 74.5±4.3 | 210.1±13.4 | 9.30±0.21 | 52.9 | 72.6±3.0 | 204.5±9.4 | 9.19±0.12 | 52.8 |
| CUAGΨGAG[f]<br>GAΨΨGΨUC | 71.2±5.6 | 207.9±18.4 | 6.74±0.13 | 37.9 | 70.4±3.5 | 205.5±11.3 | 6.70±0.05 | 37.2 |
| CUCGUGAG[f]<br>GAGΨGCUC | 44.2±3.7 | 121.0±11.7 | 6.67±0.07 | 44.1 | 50.8±2.1 | 142.0±6.7 | 6.80±0.05 | 44.0 |
| CUCGΨGAG[f]<br>GAGΨGCUC | 73.1±4.4 | 207.0±13.6 | 8.92±0.14 | 51.4 | 76.1±2.8 | 216.3±8.6 | 9.00±0.10 | 51.2 |
| CCAGΨΨGG[c,f]<br>GGΨΨGACC | 76.0±5.5 | 212.7±17.2 | 10.07±0.19 | 56.0 | 84.3±4.8 | 238.3±14.8 | 10.42±0.24 | 55.5 |
| CΨGΨAΨAG[f]<br>GAΨAΨGΨC | 58.6±2.4 | 164.9±7.7 | 7.43±0.07 | 46.6 | 61.1±1.6 | 173.1±5.0 | 7.45±0.03 | 46.3 |
| CΨGUAΨAG[f]<br>GAΨAUGΨC | 59.3±2.5 | 173.6±8.4 | 5.48±0.09 | 36.0 | 59.2±1.9 | 173.4±6.3 | 5.47±0.03 | 35.9 |
| CGΨUGUAG[f]<br>GCAGΨGΨC | 67.1±2.7 | 195.7±9.4 | 6.42±0.10 | 36.5 | 71.0±3.1 | 208.4±10.0 | 6.37±0.05 | 36.3 |
| CΨACGΨAG[f]<br>GAΨGCAΨC | 77.6±2.6 | 211.7±7.8 | 11.89±0.24 | 64.0 | 80.9±6.0 | 221.7±17.9 | 12.12±0.41 | 63.8 |
| CΨAΨAΨAG[f]<br>GAΨAΨAΨC | 70.9±3.0 | 198.0±9.2 | 9.53±0.18 | 54.8 | 77.8±3.7 | 219.1±11.4 | 9.53±0.17 | 54.5 |
| CΨGCΨAGC[c,f]<br>GAUGAΨΨG | 47.3±6.6 | 133.4±22.2 | 5.90±0.40 | 32.9 | 78.9±6.4 | 237.8±22.1 | 5.17±0.17 | 31.8 |
| CUCGGCUC[c,f]<br>GAGΨΨGAG | 76.2±4.6 | 213.6±14.4 | 9.97±0.20 | 51.7 | 71.4±3.0 | 198.6±8.8 | 9.76±0.10 | 51.7 |
| CUCGGCUC[c,f]<br>GAGΨUGAG | 74.7±3.1 | 212.0±9.6 | 8.95±0.07 | 47.4 | 74.1±1.0 | 213.4±3.0 | 8.94±0.02 | 47.3 |
| CUCGGCUC[c,f]<br>GAGUΨGAG | 76.8±7.8 | 217.2±24.3 | 9.47±0.25 | 49.3 | 74.2±2.7 | 209.2±8.3 | 9.33±0.08 | 49.2 |
| GAGGΨGAG[c,f]<br>CUCΨGCUC | 77.8±3.1 | 217.0±9.5 | 10.53±0.18 | 53.8 | 76.1±2.8 | 211.8±8.8 | 10.43±0.12 | 53.7 |
| GAGGUGAG[c,f]<br>CUCΨGCUC | 75.5±5.6 | 214.5±17.7 | 8.97±0.11 | 47.4 | 82.4±2.4 | 236.3±7.5 | 9.11±0.04 | 47.0 |
| GAGΨGGAG[c,f]<br>CUCGΨCUC | 88.3±8.9 | 246.7±27.1 | 11.73±0.42 | 56.4 | 94.0±5.0 | 264.2±15.3 | 12.04±0.24 | 56.3 |
| GAGΨGGAG[c,f]<br>CUCGUCUC | 103.2±3.4 | 293.0±10.3 | 12.31±0.25 | 55.4 | 90.8±2.0 | 255.2±6.3 | 11.69±0.10 | 55.7 |
| GAAΨGAAG[f]<br>CUΨGΨUUC | 71.2±4.4 | 208.2±14.5 | 6.67±0.15 | 37.6 | 79.0±5.3 | 233.6±7.3 | 6.58±0.09 | 37.2 |
| GAAUGAAG[f]<br>CUΨGΨUUC | 66.6±4.0 | 195.3±13.0 | 6.07±0.12 | 34.9 | 73.1±1.5 | 216.7±4.9 | 5.93±0.03 | 34.5 |

| | | | | | | | | |
|---|---|---|---|---|---|---|---|---|
| GAAUGAAG[e,f]<br>CUΨGΨUUC | 77.5±5.6 | 229.8±18.0 | 6.22±0.11 | 35.8 | 65.0±2.2 | 189.1±7.3 | 6.33±0.04 | 36.1 |
| GAAΨΨAAG[f]<br>CUUGGUUC | 61.6±3.3 | 181.6±11.2 | 5.28±0.17 | 30.8 | 63.7±3.3 | 188.5±11.1 | 5.21±0.11 | 30.7 |
| GAAGUAAG[f]<br>CUUΨGUUC | 66.2±5.5 | 200.9±18.5 | 4.00±0.28 | 25.6 | 70.2±4.0 | 214.2±13.3 | 3.78±0.18 | 25.3 |
| GAAΨΨAAG[f]<br>CΨUAGUUC | 64.1±3.2 | 186.5±9.9 | 6.30±0.24 | 35.9 | 60.5±8.0 | 174.6±25.5 | 6.33±0.24 | 36 |
| GAGΨUAAG[f]<br>CΨUAGUUC | 68.1±3.5 | 203.6±11.9 | 4.99±0.16 | 30.2 | 75.3±3.0 | 227.3±10.0 | 4.77±0.09 | 29.9 |
| GAAUGAAG[f]<br>CUUGCΨΨC | 66.7±3.2 | 192.4±9.9 | 7.05±0.18 | 39.4 | 69.4±6.2 | 201.2±19.9 | 7.03±0.11 | 39.2 |
| GCAAΨΨGC[g]<br>CGΨΨAACG | 73.4 ±7.2 | 201.1 ± 22.1 | 11.03 ± 0.41 | 61.4 | 74.1±4.4 | 203.4±13.4 | 11.03±0.27 | 61.1 |
| GΨΨCGAAC[g]<br>CAAGCΨΨG | 75.3 ±4.1 | 207.7 ± 12.4 | 10.86 ± 0.25 | 60.0 | 77.6±4.8 | 214.9±14.7 | 10.96± 0.28 | 59.7 |
| GCGAΨUGC[g]<br>CGUΨAGCG | 67.6 ±5.2 | 193.8 ± 16.6 | 7.45 ± 0.09 | 45.3 | 68.1±2.3 | 195.7±7.2 | 7.40± 0.03 | 45.1 |
| GCGGΨΨGC[g]<br>CGΨΨGGCG | 67.8 ±5.7 | 192.2 ± 18.3 | 8.17 ± 0.10 | 48.9 | 68.7±1.5 | 195.1±4.6 | 8.15± 0.03 | 48.6 |
| GΨΨCGGGC[g]<br>CGGGCΨΨG | 74.0 ±4.7 | 212.2 ±14.7 | 8.18 ± 0.15 | 47.9 | 70.0± 2.5 | 199.8±7.9 | 8.04±0.06 | 47.9 |
| GCGGΨUGC[g]<br>CGUΨGGCG | 71.9 ±3.7 | 208.5 ± 11.7 | 7.23 ± 0.07 | 43.9 | 69.6± 1.6 | 201.2±5.3 | 7.19±0.02 | 43.9 |
| GACUAGΨU[g]<br>UΨGAUCAG | 66.3 ± 4.8 | 187.5 ± 15.2 | 8.18 ± 0.10 | 49.2 | 69.8±1.9 | 198.7±6.0 | 8.21±0.05 | 48.7 |
| GUΨCGAGC[g]<br>CGAGCΨUG | 62.4 ±3.4 | 177.8 ± 11.0 | 7.24 ± 0.10 | 45.0 | 68.9±3.7 | 198.5± 11.9 | 7.29±0.07 | 44.5 |
| GΨΨAΨGGC[e,f]<br>CAAΨAΨUG | 43.3±2.9 | 114.3±9.6 | 7.84±0.36 | 46.7 | 87.6±4.8 | 256.8±24.6 | 7.90±0.12 | 41.9 |
| GΨΨAΨGGC[f]<br>CGGΨAΨΨG | 53.1±12.7 | 156.8±42.0 | 4.52±0.48 | 30.4 | 49.8±6.1 | 146.2±20.3 | 4.47±0.26 | 29.7 |
| GACGCGΨΨ[c,f]<br>ΨΨGCGCAG | 77.3±7.8 | 211.0±23.7 | 11.83±0.41 | 63.8 | 76.2±3.9 | 207.7±11.6 | 11.82±0.28 | 64.2 |
| GGΨUGACC[c,f]<br>CCAGUΨGG | 80.5±4.7 | 226.4±14.9 | 10.26±0.24 | 55.7 | 77.1±4.5 | 216.1±13.7 | 10.06±0.22 | 55.7 |
| GCUGGΨG[c,f]<br>CGAUΨACG | 74.8±4.3 | 208.1±13.9 | 10.26±0.10 | 53.3 | 80.6±10.8 | 226.1±33.7 | 10.44±0.41 | 52.8 |
| GCUGGΨGC[c,f]<br>CGAUUACG | 66.5±7.7 | 184.6±24.1 | 9.23±0.25 | 50.1 | 60.0±5.7 | 183.2±17.6 | 9.20±0.28 | 50.0 |
| GCUGGUGC[c,f]<br>CGAUΨACG | 79.6±4.6 | 227.3±14.0 | 9.09±0.27 | 47.3 | 77.2±6.4 | 219.8±20.1 | 9.02±0.21 | 47.3 |
| GCAGΨUGC[c,f]<br>CGUΨGACG | 75.7±5.0 | 214.6±15.6 | 9.14±0.26 | 51.9 | 77.1±6.8 | 219.2±21.1 | 9.16±0.30 | 51.7 |
| GCAGCUGΨ[c,f]<br>ΨGUCGACG | 79.0±2.5 | 216.4±7.8 | 11.84±0.16 | 63.2 | 84.1±6.3 | 231.8±18.9 | 12.20±0.48 | 63.1 |
| GCUGGUGC[c,f]<br>CGAΨΨACG | 70.0±3.6 | 195.5±11.2 | 9.38±0.19 | 50.1 | 61.3±4.2 | 168.2±13.1 | 9.12±0.13 | 50.7 |
| GCUGGUGC[c,f]<br>CGAΨUACG | 78.9±2.5 | 227.1±7.7 | 8.50±0.18 | 44.9 | 69.0±5.6 | 195.4±17.9 | 8.39±0.12 | 45.6 |

| | | | | | | | | |
|---|---|---|---|---|---|---|---|---|
| GGAΨGUCC[c,f]<br>CCUGΨAGG | 83.6±4.4 | 234.2±13.4 | 10.93±0.28 | 57.8 | 81.7±2.9 | 228.7±8.8 | 10.80±0.16 | 57.8 |
| GGCΨΨCAA[c,f]<br>CCGAAGUU | 69.1±3.6 | 185.6±11.4 | 11.53±0.14 | 61.2 | 73.5±3.4 | 199.0±10.3 | 11.76±0.21 | 60.8 |
| GGΨΨAΨGG[f]<br>CCAGΨGΨU | 59.0±5.5 | 165.7±17.8 | 7.66±0.09 | 43.0 | 60.3±3.8 | 169.6±12.4 | 7.70±0.08 | 43.2 |
| ΨACUAGΨA[g]<br>AΨGAUCAΨ | 70.4 ±5.3 | 196.9 ± 16.5 | 9.33 ± 0.20 | 54.0 | 74.1±2.6 | 208.6±7.9 | 9.42±0.10 | 53.5 |
| ΨGCUAGΨG[g]<br>GΨGAUCGΨ | 74.2 ± 5.5 | 208.7 ± 17.4 | 9.43 ± 0.12 | 53.5 | 75.2± 4.6 | 212.0±14.2 | 9.45±0.20 | 53.4 |
| UΨACGUAG[c,f]<br>GAUGCAΨU | 56.9±2.2 | 159.7±6.9 | 7.38±0.13 | 46.6 | 57.6±3.2 | 161.9±10.0 | 7.37±0.07 | 46.4 |
| CGΨΨAΨACG[f]<br>GCAAΨAΨGC | 84.5±2.3 | 229.1±7.0 | 13.47±0.17 | 64.7 | 79.0±2.4 | 212.7±7.1 | 13.04±0.19 | 64.8 |
| GCΨΨCAGCG[f]<br>CGAAGΨΨGC | 90.0±1.7 | 245.1±4.8 | 14.00±0.22 | 65.1 | 87.7±3.9 | 238.2±11.7 | 13.82±0.30 | 65.1 |
| GCAACΨΨCG[f]<br>CGΨΨGAAGC | 87.7±3.0 | 236.8±8.8 | 14.27±0.25 | 67.0 | 87.5±2.5 | 236.2±6.9 | 14.22±0.19 | 66.9 |
| ΨCAGUCAGU [c,d]<br>AGUCAGUCA | 76.0 ± 3.4 | 207.3 ± 10.6 | 11.64 ± 0.13 | 59.9 | 84.3 ± 2.0 | 233.0 ± 6.1 | 12.07±0.11 | 58.8 |
| ΨCAGUCAGU[c,d]<br>GGUCAGUCA | 77.7 ± 4.5 | 213.5 ± 13.7 | 11.51 ± 0.28 | 58.2 | 83.5 ± 2.4 | 231.3 ± 7.3 | 11.79 ± 0.12 | 57.8 |
| UCAGΨCAGU[c,d]<br>AGUCAGUCA | 85.0 ± 7.2 | 233.1 ± 21.9 | 12.73 ± 0.37 | 61.4 | 87.2 ± 7.0 | 239.9 ± 21.1 | 12.85 ± 0.43 | 61.2 |
| UCACΨGAGU[c,d]<br>AGUGACUCA | 105.5 ± 8.2 | 293.3 ± 24.4 | 14.51 ± 0.32 | 62.4 | 104.9 ± 6.2 | 291.6 ± 18.5 | 14.44±0.47 | 62.3 |
| UCAAΨUAGU [c,d]<br>AGUUAAUCA | 78.9 ± 7.3 | 226.8 ± 22.9 | 8.51 ± 0.18 | 45.0 | 71.5 ± 2.7 | 203.5 ± 8.7 | 8.36±0.05 | 45.2 |
| UCAUΨAAGU [c,d]<br>AGUAAUUCA | 76.7 ± 8.0 | 220.1 ± 25.1 | 8.48 ± 0.19 | 45.1 | 74.1 ± 4.1 | 211.8 ± 13.3 | 8.42±0.08 | 45.1 |
| UCAGUCAGΨ [c,d]<br>AGUCAGUCA | 79.8 ± 10.2 | 218.7 ± 31.1 | 11.97 ± 0.55 | 59.7 | 82.6 ± 6.8 | 227.3 ± 20.9 | 2.06±0.42 | 59.3 |
| UCAGΨCAGU[c,d]<br>AGUCGGUCA | 95.8 ± 10.3 | 264.9 ± 30.8 | 13.62 ± 0.76 | 61.8 | 93.4 ± 4.2 | 257.9 ± 12.8 | 13.41 ± 0.27 | 61.7 |
| UCACΨGAGU[c,d]<br>AGUGGCUCA | 82.2 ± 6.6 | 227.2 ± 19.9 | 11.77 ± 0.42 | 58.1 | 79.5 ± 2.3 | 218.8 ± 7.3 | 11.61 ± 0.12 | 58.2 |
| UCAAΨUAGU[c,d]<br>AGUUGAUCA | 75.6 ± 5.7 | 218.4 ± 18.4 | 7.89 ± 0.14 | 42.7 | 68.4 ± 5.0 | 195.5 ± 15.9 | 7.82 ± 0.10 | 43.0 |
| UCAUΨAAGU[c,d]<br>AGUAGUUCA | 78.8 ± 8.5 | 230.2 ± 27.2 | 7.41 ± 0.12 | 40.5 | 70.7 ± 2.5 | 204.0 ± 8.2 | 7.40 ± 0.02 | 40.9 |
| UCAGUCAGΨ[c,d]<br>AGUCAGUCG | 99.7 ± 5.6 | 277.6 ± 17.1 | 13.60 ± 0.35 | 60.7 | 93.1 ± 2.6 | 257.6 ± 7.9 | 13.18 ± 0.16 | 60.9 |
| CCAGCGUCCU[c,f]<br>GGΨΨGΨAGGA | 95.2±7.0 | 258.5±20.8 | 14.99±0.54 | 67.3 | 92.9±2.0 | 251.9±5.8 | 14.77±0.16 | 67.2 |
| CCAGCGUCCU[c,f]<br>GGΨUGΨAGGA | 100.4±3.8 | 277.1±11.2 | 14.43±0.33 | 63.5 | 104.0±7.6 | 287.9±22.9 | 14.68±0.54 | 63.4 |
| CΨGUCGAUAG[g]<br>GAUAGCUGΨC | 82.4 ± 3.3 | 235.9 ±10.6 | 9.28 ± 0.07 | 51.2 | 85.3±2.2 | 244.8±7.0 | 9.35±0.07 | 51.0 |

| | | | | | | | | |
|---|---|---|---|---|---|---|---|---|
| CAGAGGAGAC[c,f]<br>GUCUUΨUCUG | 113.4±10.9 | 327.5±33.5 | 11.84±0.56 | 52.2 | 101.7±7.5 | 291.2±23.0 | 11.39±0.31 | 52.5 |
| GAGΨGGAGAG[c,f]<br>CUCAΨUUCUC | 91.3±4.5 | 257.7±13.9 | 11.40±0.22 | 54.4 | 94.5±4.3 | 267.4±13.4 | 11.52±0.20 | 54.3 |
| GAGΨGGAGAG[e,f]<br>CUCAUUUCUC | 65.4±3.6 | 191.1±11.4 | 6.18±0.09 | 35.4 | 76.7±2.1 | 227.7±7.0 | 6.05±0.03 | 35.1 |
| GΨGAAUUUAC[c,f]<br>CAUUUAAGΨG | 76.7±3.6 | 229.5±11.9 | 5.48±0.13 | 36.2 | 83.0±4.2 | 250.1±13.7 | 5.42±0.07 | 36.0 |
| CΨGGUGΨΨAU[f]<br>GGΨΨAUAGΨG | 64.0±5.1 | 186.2±16.3 | 6.29±0.13 | 35.8 | 69.7±8.0 | 204.4±25.8 | 6.33±0.16 | 36.1 |
| GΨGAGCUUAC[g]<br>CAUUCGAGΨG | 81.4 ± 4.7 | 231.6 ± 14.7 | 9.54 ± 0.14 | 52.4 | 84.2±3.2 | 240.5±9.9 | 9.59±0.11 | 52.1 |
| GUΨGAUCAGC[g]<br>CGACUAGΨUG | 75.2 ± 3.8 | 210.4 ± 11.8 | 9.96 ± 0.14 | 55.7 | 82.6±3.1 | 233.2±9.7 | 10.24±0.13 | 55.1 |
| GAGΨGCAUUC[g]<br>CUUACGΨGAG | 87.2 ± 2.8 | 243.7 ± 8.5 | 11.59 ± 0.16 | 59.6 | 97.2±4.5 | 274.3±13.8 | 12.10±0.24 | 58.9 |

[a]The duplex was excluded because its residual was identified as an outlier during the linear regression analysis
[b]Calculated for $10^{-4}$ M oligomer concentration
[c]Used to compare the ΔH° and ΔS° values with analogous U containing sequences reported in [37,39,48]
[d]Reported previously [67]
[e] These duplexes were excluded from the fit because the ΔH˚ values as fit by the two methods differed by greater than 15%. This can indicate non-two-state melting
[f]Institute of Bioorganic Chemistry of Polish Academy of Sciences
[g]Saint Louis University

**Table S3.** The helical stack nearest neighbor parameters for Ψ-A pairs. σ is the uncertainty, which is the standard error of the regression. The U analogue values are from Xia et al. [48]. For each stack, the top sequence is oriented in the 5′ to 3′ direction, and the bottom sequence is written 3′ to 5′.

| **Ψ-A stack** | **$\Delta G°_{37}$ (kcal/mol)** | **σ** | **<u>U</u> analog** | **$\Delta G°_{37}$ (kcal/mol)** | **σ** | **# Occurrences in dataset[a]** | **$\Delta\Delta G°_{37}$ (Ψ stack – U analogue)** |
|---|---|---|---|---|---|---|---|
| AC<br>ΨG | -3.10 | 0.23 | AC<br>UG | -2.24 | 0.06 | 34 | -0.86±0.24 |
| AG<br>ΨC | -2.69 | 0.23 | AG<br>UC | -2.08 | 0.06 | 30 | -0.61±0.24 |
| GA<br>CΨ | -2.44 | 0.25 | GA<br>CU | -2.35 | 0.06 | 21 | -0.09±0.26 |
| CA<br>GΨ | -2.43 | 0.25 | CA<br>GU | -2.11 | 0.07 | 25 | -0.32±0.26 |
| AΨ<br>UA | -2.39 | 0.33 | AU<br>UA | -1.10 | 0.08 | 7 | -1.29±0.34 |
| ΨA<br>AU | -2.13 | 0.29 | UA<br>AU | -1.33 | 0.09 | 11 | -0.80±0.30 |
| AA<br>UΨ | -2.10 | 0.33 | AA<br>UU | -0.93 | 0.03 | 8 | -1.17±0.33 |
| AΨ<br>ΨA | -1.94 | 0.44 | AU<br>UA | -1.10 | 0.08 | 13 | -0.84±0.44 |
| ΨA<br>AΨ | -1.53 | 0.42 | UA<br>AU | -1.33 | 0.09 | 19 | -0.20±0.43 |
| AA<br>ΨU | -1.38 | 0.28 | AA<br>UU | -0.93 | 0.03 | 9 | -0.45±0.28 |
| AA<br>ΨΨ | -0.92 | 0.19 | AA<br>UU | -0.93 | 0.03 | 15 | 0.01±0.19 |
| AΨ end | 0.42 | 0.21 | AU end | 0.45 | 0.04 | 13 | -0.03±0.21 |

[a]The number of times the nearest neighbor pair appears in the duplexes that were studied by optical melting experiments (Table S1).

**Table S4.** The helical stack nearest neighbor parameters for Ψ-G pairs. σ is the uncertainty, which is the standard error of the regression. The U analogue values are from Chen et al. [37]. For each pair, the top sequence is oriented in the 5′ to 3′ direction, and the bottom sequence is written 3′ to 5′.

| **Ψ-G stacks** | **$\Delta G°_{37}$ (kcal/mol)** | **σ** | **U analog** | **$\Delta G°_{37}$ (kcal/mol)** | **σ** | **# Occurrences in dataset[a]** | **$\Delta\Delta G°_{37}$ (Ψ stack – U analogue)** |
|---|---|---|---|---|---|---|---|
| GΨ<br>CG | -3.21 | 0.20 | GU<br>CG | -2.15 | 0.10 | 22 | -1.06±0.22 |
| CΨ<br>GG | -2.76 | 0.23 | CU<br>GG | -1.77 | 0.09 | 10 | -0.99±0.25 |
| AΨ<br>ΨG | -2.33 | 0.42 | AU<br>UG | -0.90 | 0.08 | 5 | -1.43±0.43 |
| GΨ<br>ΨG | -2.28 | 0.41 | GU<br>UG | 0.72 | 0.19 | 9 | -3.00±0.45 |
| AΨ<br>UG | -1.83 | 0.22 | AU<br>UG | -0.90 | 0.08 | 16 | -0.93±0.23 |
| GG<br>CΨ | -1.79 | 0.25 | GG<br>CU | -1.80 | 0.09 | 10 | 0.01±0.27 |
| AU<br>ΨG | -1.43 | 0.32 | AU<br>UG | -0.90 | 0.08 | 7 | -0.53±0.33 |
| CG<br>GΨ | -1.39 | 0.19 | CG<br>GU | -1.25 | 0.09 | 22 | -0.14±0.21 |
| GA<br>ΨΨ | -1.19 | 0.38 | GA<br>UU | -0.51 | 0.08 | 5 | -0.68±0.39 |
| ΨG<br>GU | -1.13 | 0.49 | UG<br>GU | -0.57 | 0.19 | 3 | -0.56±0.53 |
| GA<br>ΨU | -0.97 | 0.29 | GA<br>UU | -0.51 | 0.08 | 9 | -0.46±0.30 |
| ΨG<br>GΨ | -0.97 | 0.33 | UG<br>GU | -0.57 | 0.19 | 8 | -0.40±0.38 |
| GG<br>ΨU | -0.66 | 0.42 | GG<br>UU | -0.25 | 0.16 | 4 | -0.41±0.45 |
| GU<br>ΨG | -0.62 | 0.41 | GU<br>UG | 0.72 | 0.19 | 4 | -1.34±0.45 |
| GA<br>UΨ | -0.59 | 0.29 | GA<br>UU | -0.51 | 0.08 | 9 | -0.08±0.30 |

| | | | | | | | |
|---|---|---|---|---|---|---|---|
| UG<br>AΨ | -0.41 | 0.24 | UA<br>GU | -0.39 | 0.09 | 12 | -0.02±0.26 |
| AG<br>ΨΨ | -0.40 | 0.42 | AG<br>UU | -0.35 | 0.08 | 5 | -0.05±0.43 |
| GG<br>ΨΨ | -0.30 | 0.22 | GG<br>UU | -0.25 | 0.16 | 11 | -0.05±0.27 |
| AG<br>ΨU | -0.23 | 0.29 | AG<br>UU | -0.35 | 0.08 | 8 | 0.12±0.30 |
| GG<br>UΨ | -0.20 | 0.30 | GG<br>UU | -0.25 | 0.16 | 6 | 0.05±0.34 |
| AG<br>UΨ | -0.18 | 0.26 | AG<br>UU | -0.35 | 0.08 | 9 | 0.17±0.27 |
| ΨG<br>AΨ | -0.06 | 0.35 | UG<br>AU | -0.39 | 0.09 | 9 | 0.33±0.36 |
| UA<br>GΨ | 0.18 | 0.29 | UG<br>AU | -0.39 | 0.09 | 8 | 0.57±0.30 |
| GΨ<br>end | 0.21 | 0.15 | GU end | 0.00 | 0.00 | 21 | 0.21±0.15 |

[a]The number of times the nearest neighbor pair appears in the duplexes that were studied by optical melting experiments

**Table S5.** Optical melting experiment results for duplexes with dangling ends. ΔH° is the folding enthalpy change, ΔS° is the folding entropy change, and $\Delta G°_{37}$ is the folding free energy change at 37 °C, determined from the enthalpy and entropy changes. $T_M$ is the melting temperature. Pseudouridines and nucleotides paired with pseudouridine are are shown in red, and the nearest neighbor to the pseudouridines are shown in blue.

| Duplexes (5'-3') | Average of curve fits | | | | $T_M^{-1}$ vs log $C_T$ plots | | | |
|---|---|---|---|---|---|---|---|---|
| | -ΔH° (kcal/mol) | -ΔS° (eu) | $-\Delta G°_{37}$ (kcal/mol) | $T_M$[b] (°C) | -ΔH° (kcal/mol) | -ΔS° (eu) | $-\Delta G°_{37}$ (kcal/mol) | $T_M$[b] (°C) |
| ΨAUGCAU[d]<br>UACGUAΨ | 45.5±4.4 | 128.9±13.9 | 5.51±0.09 | 35.9 | 49.7±2.9 | 142.5±9.6 | 5.50±0.05 | 35.9 |
| AUGCAUΨ[d]<br>ΨUACGUA | 46.2±1.5 | 130.0±4.7 | 5.91±0.11 | 38.6 | 53.0±2.7 | 152.2±8.9 | 5.84±0.05 | 38.0 |
| AUGCAΨΨ[a,d]<br>ΨΨACGUA | 43.4±1.8 | 121.0±5.4 | 5.92±0.12 | 38.7 | 41.4±3.3 | 114.5±10.6 | 5.93±0.08 | 38.9 |
| AUGCAΨG[d]<br>GΨACGUA | 51.3±1.6 | 142.9±5.1 | 6.97±0.07 | 45.0 | 56.5±1.7 | 159.6±5.3 | 7.05±0.03 | 44.7 |
| CΨCAUGA[d]<br>AGUACΨC | 50.2±2.7 | 143.5±8.9 | 5.65±0.12 | 36.9 | 53.4±3.2 | 154.0±10.4 | 5.64±0.07 | 36.8 |
| ΨGCGCAA[d]<br>AACGCGΨ | 62.0±3.6 | 165.6±10.5 | 10.60±0.34 | 63.8 | 63.8±5.7 | 168.3±17.0 | 10.70±0.41 | 63.9 |
| ACAUGUΨ[e]<br>ΨUGUACA | 53.0±2.8 | 154.4±9.5 | 5.16±0.15 | 34.0 | 53.2±2.0 | 155.1±6.5 | 5.15±0.05 | 34.0 |
| GCGCΨ[e]<br>ΨCGCG | 54.4±6.1 | 151.0±19.7 | 7.54±0.15 | 48.0 | 55.5±5.1 | 154.6±16.1 | 7.53±0.17 | 47.8 |
| CCGGΨ[e]<br>ΨGGCC | 47.8±4.1 | 133.9±13.0 | 6.30±0.07 | 41.1 | 46.8±1.8 | 130.7±5.9 | 6.26±0.03 | 40.9 |
| ΨAUGCAU[e]<br>_ UACGUAΨ | 49.1±5.6 | 140.2±18.2 | 5.60±0.14 | 36.5 | 45.7±4.6 | 129.3±14.9 | 5.60±0.15 | 36.5 |
| ΨCAGUCAGU[c]<br>_ GUCAGUCA | 78.5±2.8 | 216.8±8.5 | 11.23±0.14 | 56.8 | 71.7±1.4 | 195.8±4.3 | 10.92±0.06 | 57.2 |
| UCAGUCAGΨ[c]<br>AGUCAGUC | 77.1±8.3 | 212.7±25.5 | 11.17±0.44 | 56.9 | 72.9±5.1 | 199.9±15.8 | 10.93±0.25 | 56.9 |

[a] Excluded from additional analysis because the duplex has a dangling end that is too destabilizing.

[b] Calculated for $10^{-4}$ M oligomer concentration

[c] Reported previously[37]

[d] Institute of Bioorganic Chemistry of Polish Academy of Sciences

[e] Saint Louis University

**Table S6.** Dangling end motif stability nearest neighbor parameters for both 5′ and 3′ dangling ends. Source indicates if the $\Delta G^\circ_{37}$ is measured or estimated.

| Motif | Source | $\Delta G^\circ_{37}$ (kcal/mol) | $\Delta G^\circ_{37}$ U analog (kcal/mol) | $\Delta\Delta G^\circ_{37\,\text{dangle}}$[a] (kcal/mol) |
|---|---|---|---|---|
| ΨA<br>U | Estimate | -0.64±0.16 | -0.20±0.14 | -0.44±0.07[b] |
| ΨC<br>G | Estimate | -0.51±0.11 | -0.07±0.09 | -0.44±0.07[b] |
| ΨG<br>C | Experiment | -0.66±0.30 | -0.15±0.12 | -0.51±0.32 |
| ΨU<br>A | Experiment | -0.57±0.14 | -0.19±0.11 | -0.38±0.18 |
| ΨG<br>U | Estimate | -0.64±0.16 | -0.20±0.14 | -0.44±0.07[b] |
| ΨU<br>G | Estimate | -0.63±0.13 | -0.19±0.11 | -0.44±0.07[b] |
| AA<br>Ψ | Estimate | -0.56±0.89 | -0.30±0.13 | -0.26±0.39 |
| CA<br>Ψ | Experiment | -0.40±0.37 | -0.14±0.11 | -0.26±0.39 |
| GA<br>Ψ | Estimate | -0.46±0.41 | -0.20±0.14 | -0.26±0.39 |
| UA<br>Ψ | Estimate | -0.46±0.41 | -0.20±0.14 | -0.26±0.39 |
| ΨA<br>Ψ | Estimate | -0.64±0.16 | -0.20±0.14 | -0.44±0.07[b] |
| AΨ<br>A | Estimate | -0.59±0.41 | -0.33±0.12 | -0.26±0.39 |
| CΨ<br>A | Estimate | -0.51±0.40 | -0.25±0.10 | -0.26±0.39 |
| GΨ<br>A | Estimate | -0.61±0.41 | -0.35±0.12 | -0.26±0.39 |
| UΨ<br>A | Estimate | -0.45±0.40 | -0.19±0.11 | -0.26±0.39 |
| ΨΨ<br>A | Estimate | -0.63±0.13 | -0.19±0.11 | -0.44±0.07[b] |
| AΨ<br>U | Estimate | -0.92±0.31 | -0.58±0.30 | -0.34±0.06[b] |

| | | | | |
|---|---|---|---|---|
| CΨ<br>G | Experiment | -1.46±0.18 | -1.19±0.12 | -0.27±0.21 |
| GΨ<br>C | Experiment | -0.91±0.17 | -0.62±0.13 | -0.30±0.21 |
| UΨ<br>A | Experiment | -0.55±0.14 | -0.09±0.17 | -0.46±0.22 |
| GΨ<br>U | Estimate | -0.92±0.31 | -0.58±0.30 | -0.34±0.06 [b] |
| UΨ<br>G | Estimate | -0.43±0.18 | -0.09±0.17 | -0.34±0.06 [b] |
| AA<br>Ψ | Experiment | -1.0±0.42 | -0.74±0.29 | -0.26±0.51 |
| AC<br>Ψ | Estimate | -0.75±0.53 | -0.49±0.15 | -0.26±0.51 |
| AG<br>Ψ | Estimate | -1.09±0.58 | -0.83±0.29 | -0.26±0.51 |
| AU<br>Ψ | Estimate | -0.84±0.59 | -0.58±0.30 | -0.26±0.51 |
| AΨ<br>Ψ | Estimate | -0.92±0.31 | -0.58±0.30 | -0.34±0.06 [b] |
| ΨA<br>A | Estimate | 0.00±0.48 | -0.66±0.16 | 0.67±0.46 |
| ΨC<br>A | Estimate | 0.54±0.47 | -0.13±0.13 | 0.67±0.46 |
| ΨG<br>A | Experiment | 0.00±0.44 | -0.66±0.13 | 0.67±0.46 |
| ΨU<br>A | Estimate | 0.58±0.49 | -0.09±0.17 | 0.67±0.46 |
| ΨΨ<br>A | Estimate | -0.43±0.18 | -0.09±0.17 | -0.34±0.06 |

[a]For values that are from an experimental source, the uncertainty was propogated from the uncertainties of $\Delta G^\circ_{37}$ and the $\Delta G^\circ_{37}$ for the U analog. For estimated values, the uncertainity is of the $\Delta\Delta G^\circ_{37\ dangle}$ that was used to estimate the $\Delta G^\circ_{37}$ from a $\Delta G^\circ_{37}$ for the U analog, as described in the text, unless indicated otherwise.

[b]Uncertainty is the standard error of the mean of the $\Delta\Delta G^\circ_{37}$ values that were averaged to extrapolate the $\Delta G^\circ_{37}$ value as described in the text.

**Table S7:** Optical melting data for duplexes containing terminal mismatches. ΔH° is the folding enthalpy change, ΔS° is the folding entropy change, and $\Delta G°_{37}$ is the folding free energy change at 37 °C, determined from the enthalpy and entropy changes. $T_M$ is the melting temperature. The pseudouridine base pairs and pseudouridine terminal mismatches are shown in red, and the nearest neighbors to the pseudouridines are shown in blue.

| Duplexes (5'-3') | Average of curve fits | | | | $T_M^{-1}$ vs log $C_T$ plots | | | |
|---|---|---|---|---|---|---|---|---|
| | -ΔH° (kcal/mol) | -ΔS° (eu) | $-\Delta G°_{37}$ (kcal/mol) | $T_M$[b] (°C) | -ΔH° (kcal/mol) | -ΔS° (eu) | $-\Delta G°_{37}$ (kcal/mol) | $T_M$[b] (°C) |
| AΨGCGCAA[d]<br>AACGCGΨA | 60.6±2.4 | 160.5±7.1 | 10.79±0.18 | 65.6 | 61.3±2.0 | 162.7±6.0 | 10.86±0.15 | 65.7 |
| UCAGUCAGΨ[d]<br>AGUCAGUCΨ | 72.7±2.5 | 199.1±7.8 | 10.93±0.16 | 57.0 | 81.7±2.8 | 226.5±8.6 | 11.40±0.15 | 56.6 |
| AAUGCAΨC[c,d]<br>CΨACGUAA | 40.8±5.2 | 111.6±16.8 | 6.15±0.13 | 40.7 | 47.0±3.0 | 131.8±9.9 | 6.13±0.07 | 40.0 |
| AΨGCGCAC[d]<br>CACGCGΨA | 62.9±3.8 | 167.1±11.1 | 11.10±0.35 | 66.2 | 68.1±3.5 | 182.6±10.3 | 11.50±0.26 | 66.0 |
| CAUGCAΨA[d]<br>AΨACGUAC | 49.5±2.0 | 136.8±6.6 | 7.11±0.09 | 46.2 | 55.7±4.6 | 156.3±14.7 | 7.23±0.14 | 45.9 |
| ΨGCGCΨ[d]<br>ΨCGCGΨ | 51.8±4.9 | 138.9±15.2 | 8.71±0.22 | 56.3 | 58.1±2.9 | 158.6±9.0 | 8.97±0.13 | 55.6 |
| ΨΨGCGCGΨ[e]<br>ΨGCGCGΨΨ | 56.3±2.2 | 150.3±6.8 | 9.70±0.12 | 60.8 | 58.5±2.1 | 157.1±6.3 | 9.80±0.11 | 60.5 |
| ΨΨGCGCAΨ[e]<br>ΨACGCGΨΨ | 65.6±2.2 | 175.0±6.8 | 11.29±0.09 | 66.0 | 66.0±3.4 | 176.3±10.3 | 11.32±0.24 | 66.0 |
| ΨUGCGCAΨ[e]<br>ΨACGCGUΨ | 59.7±2.7 | 159.8±8.7 | 10.16±0.10 | 62.2 | 60.4±5.7 | 161.8±17.2 | 10.19±0.36 | 62.1 |
| ΨGCAUGCΨ[e]<br>ΨCGUACGΨ | 73.9±3.8 | 202.1±11.5 | 11.22±0.34 | 62.2 | 74.8±6.2 | 204.9±18.6 | 11.25±0.41 | 62.0 |
| ΨCAGUCAGU[a]<br>UGUCAGUCA | 87.4 ± 6.8 | 243.4 ± 20.8 | 11.95 ± 0.36 | 57.5 | 82.3 ± 1.7 | 227.7 ± 5.2 | 11.67 ± 0.09 | 57.7 |
| UCAGUCAGΨ[a]<br>AGUCAGUCU | 83.4 ± 5.1 | 231.2 ± 15.7 | 11.70 ± 0.27 | 57.5 | 85.7 ± 4.1 | 238.2 ± 12.6 | 11.79 ± 0.22 | 57.3 |
| ΨCAGUCAGU[a]<br>CGUCAGUCA | 74.4 ± 3.8 | 204.1 ± 11.9 | 11.08 ± 0.20 | 57.2 | 74.9 ± 3.4 | 206.0 ± 10.6 | 11.07 ± 0.15 | 57.0 |
| UCAGUCAGΨ[a]<br>AGUCAGUCC | 77.7 ± 3.3 | 214.1 ± 10.3 | 11.32 ± 0.16 | 57.4 | 81.6 ± 2.2 | 226.0 ± 6.8 | 11.48 ± 0.11 | 57.0 |

[a]Reported previously[67]

[b]Calculated for $10^{-4}$ M oligomer concentration.

[c] Excluded from further analysis because the duplex has a terminal mismatch that is destabilizing.

[d]Institute of Bioorganic Chemistry of the Polish Academy of Sciences.

[e]Saint Louis University.

**Table S8.** Terminal mismatch stabilities used in the nearest neighbor parameters. Source indicates whether the mismatch was measured or extrapolated as explained in the text.

| Mismatch | Source | $\Delta G^\circ_{37}$ (kcal/mol) | $\Delta G^\circ_{37}$ (U analog) (kcal/mol) | $\Delta\Delta G^\circ_{37\ mismatch}$ (kcal/mol) |
|---|---|---|---|---|
| AC<br>UΨ | Estimate | -1.02±0.66 | -0.71±0.21 | -0.31±0.62[b] |
| AU<br>UΨ | Estimate | -1.44±0.49 | -0.80±0.33 | -0.64±0.36 [b] |
| AΨ<br>UC | Estimate | -1.26±0.71 | -0.72±0.32 | -0.54±0.63 [b] |
| AΨ<br>UU | Estimate | -1.86±0.72 | -0.80±0.33 | -1.06±0.64 [b] |
| AΨ<br>UΨ | Experiment | -0.99±0.26 | -0.80±0.31 | -0.18±0.40 [a] |
| CC<br>GΨ | Estimate | -1.07±0.75 | -0.76±0.42 | -0.31±0.62 [b] |
| CU<br>GΨ | Estimate | -1.83±0.63 | -1.19±0.52 | -0.64±0.35 [b] |
| CΨ<br>GC | Estimate | -1.90±0.83 | -1.36±0.53 | -0.54±0.63 [b] |
| CΨ<br>GU | Estimate | -2.25±0.83 | -1.19±0.52 | -1.06±0.64 [b] |
| CΨ<br>GΨ | Experiment | -2.06±0.17 | -1.19±0.52 | -0.87±0.55 [a] |
| GC<br>CΨ | Experiment | -0.81±0.60 | -0.50±0.16 | -0.31±0.62 [a] |
| GU<br>CΨ | Experiment | -1.41±0.31 | -0.77±0.18 | -0.64±0.36 [a] |
| GΨ<br>CC | Experiment | -1.52±0.61 | -0.98±0.18 | -0.54±0.63 [a] |
| GΨ<br>CU | Experiment | -1.83±0.62 | -0.77±0.18 | -1.06±0.64 [a] |
| GΨ<br>CΨ | Experiment | -1.44±0.61 | -0.77±0.18 | -0.67±0.63 [a] |

| | | | | |
|---|---|---|---|---|
| UC AΨ | Estimate | -0.77±0.83 | -0.46±0.55 | -0.31±0.62 [b] |
| UU AΨ | Estimate | -1.15±0.64 | -0.51±0.53 | -0.64±0.35 [b] |
| UΨ AC | Estimate | -1.13±0.92 | -0.59±0.67 | -0.54±0.63 [b] |
| UΨ AU | Estimate | -1.57±0.83 | -0.51±0.53 | -1.06±0.64 [b] |
| UΨ AΨ | Estimate | -1.08±0.57 | -0.51±0.53 | -0.57±0.20 [e] |
| GC UΨ | Estimate | -1.02±0.65 | -0.71±0.21 | -0.31±0.68[c] |
| GU UΨ | Estimate | -1.44±0.49 | -0.57±0.54 | -0.87 ±0.72[c] |
| GΨ UC | Estimate | -1.26±0.70 | -0.72±0.32 | -0.54±0.77[c] |
| GΨ UU | Estimate | -1.86±0.72 | -0.57±0.54 | -1.29±0.90[c] |
| GΨ UΨ | Estimate | -0.99±0.26 | -0.57±0.54 | -0.41±0.60 [c] |
| UC GΨ | Estimate | -0.77±0.83 | -0.46±0.55 | -0.31±1.00 [c] |
| UU GΨ | Estimate | -1.15±0.63 | -0.40±0.22 | -0.75±0.67 [c] |
| UΨ GC | Estimate | -1.13±0.92 | -0.59±0.67 | -0.54±1.14 [c] |
| UΨ GU | Estimate | -1.57±0.83 | -0.40±0.22 | -1.17±0.86 [c] |
| UΨ GΨ | Estimate | -1.08±0.56 | -0.40±0.22 | -0.68 ±0.61 [c] |
| AA ΨA | Experiment | -1.08±0.42 | -0.75±0.42 | -0.33±0.59 [a] |
| AA ΨC | Estimate | -1.44±0.38 | -0.88±0.31 | -0.56±0.23 [e] |
| AA ΨG | Estimate | -1.33±0.32 | -0.78±0.23 | -0.56±0.23 [e] |

| | | | | |
|---|---|---|---|---|
| AC ΨA | Experiment | -1.40±0.42 | -0.62±0.41 | -0.78±0.59 [a] |
| AC ΨC | Estimate | -1.19±0.29 | -0.63±0.19 | -0.56±0.23 [e] |
| AC ΨU | Estimate | -1.27±0.32 | -0.71±0.23 | -0.56±0.23 [e] |
| AC ΨΨ | Estimate | -1.22±0.56 | -0.71±0.21 | -0.51±0.52 [b] |
| AG ΨA | Estimate | -1.36±0.37 | -0.80±0.29 | -0.56±0.23 [e] |
| AG ΨG | Estimate | -1.33±0.34 | -0.78±0.26 | -0.56±0.23 [e] |
| AU ΨC | Estimate | -1.28±0.40 | -0.72±0.32 | -0.56±0.23 [e] |
| AU ΨU | Estimate | -1.36±0.40 | -0.80±0.33 | -0.56±0.23 [e] |
| AU ΨΨ | Estimate | -1.31±0.62 | -0.80±0.33 | -0.51±0.52 [b] |
| AΨ ΨC | Estimate | -1.23±0.61 | -0.72±0.32 | -0.51±0.52 [b] |
| AΨ ΨU | Estimate | -1.31±0.62 | -0.80±0.33 | -0.51±0.52 [b] |
| AΨ ΨΨ | Experiment | -1.31±0.42 | -0.80±0.31 | -0.51±0.52 [a] |
| ΨA AA | Estimate | -0.27±0.86 | -0.94±0.52 | 0.68±0.69 [b] |
| ΨA AC | Experiment | -0.09±0.44 | -0.76±0.53 | 0.68±0.69 [a] |
| ΨA AG | Estimate | -0.43±0.81 | -1.10±0.42 | 0.68±0.69 [b] |
| ΨC AA | Estimate | -0.05±0.80 | -0.72±0.39 | 0.68±0.69 [b] |
| ΨC AC | Estimate | 0.09±0.80 | -0.59±0.42 | 0.68±0.69 [b] |
| ΨC AU | Estimate | 0.22±0.88 | -0.46±0.55 | 0.68±0.69 [b] |

| | | | | |
|---|---|---|---|---|
| ΨC<br>AΨ | Estimate | -0.10±1.08 | -0.46±0.55 | 0.36±0.69 [d] |
| ΨG<br>AA | Estimate | -0.44±0.90 | -1.11±0.58 | 0.68±0.69 [b] |
| ΨG<br>AG | Estimate | -0.48±1.00 | -1.15±0.59 | 0.68±0.69 [b] |
| ΨU<br>AC | Estimate | 0.09±0.96 | -0.59±0.67 | 0.68±0.69 [b] |
| ΨU<br>AU | Estimate | 0.17±0.87 | -0.51±0.53 | 0.68±0.69 [b] |
| ΨU<br>AΨ | Estimate | -0.48±0.94 | -0.51±0.53 | 0.03±0.69 [d] |
| ΨΨ<br>AC | Estimate | -0.46±1.15 | -0.59±0.67 | 0.14±0.69 [d] |
| ΨΨ<br>AU | Estimate | -0.90±1.08 | -0.51±0.53 | -0.38±0.69 [d] |
| ΨΨ<br>AΨ | Estimate | -0.41±0.89 | -0.51±0.53 | 0.10±0.69 [d] |
| GA<br>ΨA | Estimate | -1.08±0.42 | -0.25±0.56 | -0.83±0.70 [c] |
| GA<br>ΨC | Estimate | -1.44±0.38 | -0.88±0.31 | -0.56±0.49 [c] |
| GA<br>ΨG | Estimate | -1.33±0.32 | -0.78±0.23 | -0.55±0.39 [c] |
| GC<br>ΨA | Estimate | -1.40±0.42 | -0.62±0.41 | -0.78±0.59 [c] |
| GC<br>ΨC | Estimate | -1.19±0.30 | -0.63±0.19 | -0.56±0.36 [c] |
| GC<br>ΨU | Estimate | -1.27±0.32 | -0.71±0.23 | -0.56±0.39 [c] |
| GC<br>ΨΨ | Estimate | -1.22±0.56 | -0.71±0.21 | -0.51±0.60 [c] |
| GG<br>ΨA | Estimate | -1.36±0.37 | -0.58±0.42 | -0.78±0.55 [c] |
| GG<br>ΨG | Estimate | -1.33±0.34 | -0.78±0.26 | -0.55±0.42 [c] |

| | | | | |
|---|---|---|---|---|
| GU<br>ΨC | Estimate | -1.28±0.39 | -0.72±0.32 | -0.56±0.50 [c] |
| GU<br>ΨU | Estimate | -1.36±0.40 | -0.57±0.54 | 0.79±0.67 [c] |
| GU<br>ΨΨ | Estimate | -1.31±0.62 | -0.57±0.54 | 0.74±0.82 [c] |
| GΨ<br>UC | Estimate | -1.23±0.62 | -0.72±0.32 | -0.51±0.69 [c] |
| GΨ<br>ΨU | Estimate | -1.31±0.62 | -0.57±0.54 | 0.74±0.82 [c] |
| GΨ<br>ΨΨ | Experiment | -1.38±0.42 | -0.57±0.54 | -0.81±0.68 [a] |
| ΨA<br>GA | Estimate | -0.27±0.86 | -1.27±0.54 | 1.00±1.02 [c] |
| ΨA<br>GC | Estimate | -0.09±0.44 | -0.76±0.53 | 0.68±0.69 [c] |
| ΨA<br>GG | Estimate | -0.43±0.81 | -1.10±0.42 | 0.68±0.91 [c] |
| ΨC<br>GA | Estimate | -0.05±0.79 | -0.72±0.39 | 0.68±0.88 [c] |
| ΨC<br>GC | Estimate | 0.09±0.81 | -0.59±0.42 | 0.68±0.91 [c] |
| ΨC<br>GU | Estimate | 0.22±0.88 | -0.46±0.55 | 0.68±1.03[c] |
| ΨC<br>GΨ | Estimate | -0.10±1.08 | -0.46±0.55 | 0.36±1.21 [c] |
| ΨG<br>GA | Estimate | -0.44±0.90 | -0.48±0.56 | 0.04±1.06 [c] |
| ΨG<br>GG | Estimate | -0.48±0.91 | -0.81±0.55 | 0.33±1.06 [c] |
| ΨU<br>GC | Estimate | 0.09±0.96 | -0.59±0.67 | 0.68±1.17 [c] |
| ΨU<br>GU | Estimate | 0.17±0.87 | -0.40±0.22 | 0.57±0.90[c] |
| ΨU<br>GΨ | Estimate | -0.48±0.94 | -0.40±0.22 | -0.08±0.97 [c] |

| | | | | |
|---|---|---|---|---|
| ΨΨ<br>GC | Estimate | -0.46±1.15 | -0.59±0.67 | 0.14±1.33 [c] |
| ΨΨ<br>GU | Estimate | -0.90±1.08 | -0.40±0.22 | -0.49±1.10 [c] |
| ΨΨ<br>GΨ | Estimate | -0.41±0.89 | -0.40±0.22 | -0.01±0.92 [c] |

[a]Uncertainty was calculated from the error of $\Delta G^{\circ}_{37}$ of the mismatch that was calculated from an experiment and the U analog.

[b]Uncertainty is of the $\Delta\Delta G^{\circ}_{37\ mismatch}$ that was used to extrapolate the experiment, as described in the text.

[c]Uncertainty was calculated from the errors of the $\Delta G^{\circ}$ values extrapolated from the corresponding A–U or U–A values as described in the text, and their U analogues.

[d]Uncertainty was calculated from the errors of the of the $\Delta\Delta G^{\circ}_{37}$ values that were added to extrapolate that $\Delta\Delta G^{\circ}_{37\ mismatch}$ value as described in the text.

[e]Uncertainty is the standard error of the mean of the values that were averaged to extrapolate the $\Delta\Delta G^{\circ}_{37}$ value as described in the text.

**Table S9.** Optical melting data for duplexes containing duplexes with internal loops. ΔH° is the folding enthalpy change, ΔS° is the folding entropy change, and $\Delta G°_{37}$ is the folding free energy change at 37 °C, determined from the enthalpy and entropy changes. $T_M$ is the melting temperature. Unless indicated otherwise, these duplexes were all melted at the Institute of Bioorganic Chemistry Polish Academy of Sciences. The Internal loop is shown in red, and the closing base pairs are shown in blue.

| Sequences (5'-3') | Average of curve fits | | | | $T_M^{-1}$ vs log $C_T$ plots | | | |
|---|---|---|---|---|---|---|---|---|
| | -ΔH° (kcal/mol) | -ΔS° (eu) | $-\Delta G°_{37}$ (kcal/mol) | $T_M$[b] (°C) | -ΔH° (kcal/mol) | -ΔS° (eu) | $-\Delta G°_{37}$ (kcal/mol) | $T_M$[b] (°C) |
| UCAGΨCAGU<br>AGUCΨGUCA | 79.0±3.8 | 220.0±11.7 | 10.77±0.16 | 54.6 | 77.7±2.0 | 216.1±6.1 | 10.72±0.09 | 54.7 |
| GCAΨUCG<br>CGUUAGC | 55.2±2.1 | 158.5±6.8 | 6.03±0.09 | 34.2 | 61.1±1.9 | 178.1±6.1 | 5.92±0.04 | 33.9 |
| CGCΨGCG<br>GCGΨCGC | 61.0±2.1 | 167.6±6.4 | 9.04±0.13 | 55.1 | 59.9±1.1 | 164.3±3.4 | 8.97±0.05 | 55.0 |
| GCAΨUUGC<br>CGUUΨACG | 69.2±5.7 | 197.4±17.5 | 7.93±0.31 | 47.5 | 64.3±4.4 | 182.3±13.8 | 7.78±0.13 | 47.5 |
| GCAΨΨUGC<br>CGUΨΨACG | 72.7±5.3 | 204.4±16.2 | 9.32±0.26 | 53.4 | 68.8±2.5 | 192.3±7.7 | 9.11±0.10 | 53.3 |
| GCGUΨUCGC<br>CGCUΨUGCG | 74.4±4.1 | 213.9±13.1 | 8.02±0.11 | 47.1 | 69.1±2.8 | 197.1±8.8 | 7.92±0.05 | 47.4 |
| GCGΨUUCGC<br>CGCUUΨGCG | 79.9±7.8 | 225.5±23.9 | 10.00±0.40 | 54.7 | 75.3±1.9 | 211.6±5.8 | 9.71±0.09 | 54.6 |
| GCGΨUΨCGC<br>CGCΨUΨGCG | 84.1±3.1 | 235.3±9.4 | 11.17±0.23 | 58.7 | 84.7±10.1 | 236.0±30.8 | 11.19±0.58 | 58.6 |
| GCAUUΨGC<br>CGΨUUACG | 67.0±2.2 | 193.7±6.8 | 6.94±0.12 | 42.9 | 64.0±1.9 | 184.2±5.9 | 6.89±0.02 | 43.0 |
| GCAUΨΨGC<br>CGΨΨUACG | 72.3±5.9 | 205.7±18.4 | 8.55±0.18 | 49.8 | 72.2±2.8 | 205.3±8.8 | 8.49±0.09 | 49.8 |
| UCAGΨCAGU[a]<br>AGUCUGUCA | 77.2±7.7 | 217.9 ± 24.0 | 9.62±0.27 | 49.9 | 69.4±4.2 | 193.7±13.2 | 9.35 ± 0.13 | 50.1 |
| UCAGΨCAGU[a]<br>AGUCCGUCA | 69.6±5.3 | 197.2 ± 17.0 | 8.41±0.10 | 45.6 | 67.2±1.5 | 189.8 ± 4.7 | 8.34 ± 0.02 | 45.6 |
| CΨGCΨGG[c]<br>GGΨCGΨC | 43.0±2.9 | 118.0±9.2 | 6.43±0.08 | 42.5 | 46.3±1.2 | 128.6±3.8 | 6.42±0.01 | 42.1 |

[a]Reported previously [67].

[b]Calculated for $10^{-4}$ M oligomer concentration.

[c] This sequence was excluded from further analysis because the internal loop appeared to be too stable in relation to other 1×1 internal loops.

**Table S10.** Optical melting data for duplexes containing bulge loops. ΔH° is the folding enthalpy change, ΔS° is the folding entropy change, and $\Delta G°_{37}$ is the folding free energy change at 37 °C, determined from the enthalpy and entropy changes. $T_M$ is the melting temperature. The nucleotide in the bulge loop is shown in red and the closing base pairs are shown in blue.

| Sequences (5'-3') | Average of curve fits | | | | $T_M^{-1}$ vs log $C_T$ plots | | | |
|---|---|---|---|---|---|---|---|---|
| | -ΔH° (kcal/mol) | -ΔS° (eu) | $-\Delta G°_{37}$ (kcal/mol) | $T_M$[a] (˚C) | -ΔH° (kcal/mol) | -ΔS° (eu) | $-\Delta G°_{37}$ (kcal/mol) | $T_M$[a] (˚C) |
| GCGΨGCG[b,c]<br>CGC--CGC | 67.0±3.2 | 182.7±9.8 | 10.32±0.21 | 55.6 | 67.2±3.0 | 183.5±9.0 | 10.31±0.16 | 55.5 |
| GACCΨUGUC[c]<br>CUGG--ACAG | 60.6±2.7 | 166.3±8.9 | 9.02±0.07 | 50.3 | 69.4±1.6 | 194.0±5.0 | 9.26±0.05 | 49.7 |
| UGACΨCUCA[c]<br>ACUG--GAGU | 54.8±4.5 | 151.5±14.4 | 7.76±0.11 | 44.1 | 55.5±2.7 | 153.9±8.5 | 7.78±0.05 | 44.1 |
| CCACGΨCUCG[d]<br>GGUG--AGAGC | 79.6±4.9 | 222.2±15.4 | 10.71±0.17 | 54.2 | 82.2±3.4 | 230.3±10.6 | 10.77±0.14 | 53.9 |
| GUCCGΨCUC[d]<br>CAGG--AGAG | 73.5±4.5 | 209.2±14.4 | 8.63±0.10 | 46.1 | 73.8±3.0 | 210.3±9.4 | 8.60±0.06 | 45.9 |
| GUCAGΨCUC[d]<br>CAG--CAGAG | 59.1±6.2 | 163.7±19.8 | 8.31±0.20 | 46.6 | 57.9±4.9 | 160.3±15.5 | 8.23±0.16 | 46.4 |
| GCAUCΨ--AGGC[d]<br>UGUAGACUCCG | 95.0±7.8 | 268.2±24.0 | 11.85±0.35 | 55.4 | 95.5±4.6 | 269.7±14.2 | 11.83±0.21 | 55.2 |
| ACUGΨGAGU[d]<br>UGAC--CUCA | 61.8±5.0 | 174.9±16.1 | 7.59±0.05 | 42.4 | 60.1±1.7 | 169.5±5.5 | 7.52±0.02 | 42.2 |
| GGCGGΨCUC[d]<br>CCGCC--GAG | 65.5±7.8 | 171.5±23.3 | 12.35±0.58 | 67.2 | 66.0±9.0 | 173.0±26.8 | 12.21±0.77 | 66.8 |
| CCAAGΨCUCG[d]<br>GGUU--GGAGC | 74.0±11.5 | 211.5±37.0 | 8.40±0.11 | 45.1 | 69.5±2.6 | 197.0±8.1 | 8.36±0.04 | 45.4 |
| GUCCGΨCUC[d]<br>CAGG--GGAG | 67.4±12.7 | 191.7±40.2 | 7.92±0.31 | 43.5 | 66.4±5.1 | 189.0±16.5 | 7.8±0.11 | 43.0 |
| GUCAGΨCUC[d]<br>CAG--CGGAG | 62.1±9.0 | 174.2±28.7 | 8.04±0.14 | 44.7 | 63.0±5.4 | 177.4±17.1 | 8.01±0.15 | 44.5 |

[a]Calculated for $10^{-4}$ M oligomer concentration.
[b]This loop was excluded from analysis because the loop is much more stable than all other bulge loops.
[c]Institute of Bioorganic Chemistry of the Polish Academy of Sciences.
[d]Saint Louis University.

**Table S11:** Bulge loop stabilities. The nucleotide in the bulge loop is shown in red and the closing base pairs are shown in blue.

| Sequences (5'-3') | $\Delta G°°_{37\ loop}$ (kcal/mol) | $\Delta G°_{37\ loop}$ (U analog/Prediction) (kcal/mol) | $\Delta\Delta G°_{37\ loop}$ (kcal/mol) |
|---|---|---|---|
| GCGΨGCG[a]<br>CGC--CGC | 0.09±0.59 | 3.89±0.62 | -3.80±0.85 |
| GACCΨUGUC<br>CUGG--ACAG | 3.28±0.48 | 3.89±0.62 | -0.61±0.78 |
| UGACΨCUCA<br>ACUG--GAGU | 4.56±0.58 | 3.89±0.62 | 0.67±0.85 |
| CCACGΨCUCG<br>GGUG--AGAGC | 4.67±0.61 | 3.89±0.62 | 0.78±0.87 |
| GUCCGΨCUC<br>CAGG--AGAG | 4.72±0.56 | 3.89±0.62 | 0.83±0.83 |
| GUCAGΨCUC<br>CAG--CAGAG | 4.60±0.55 | 3.89±0.62 | 0.71±0.83 |
| GCAUCΨ--AGGC<br>UGUAGACUCCG | 5.37±0.67 | 3.89±0.62 | 1.48±0.91 |
| ACUGΨGAGU<br>UGAC--CUCA | 3.85±0.43 | 3.89±0.62 | -0.04±0.76 |
| GGCGGΨCUC<br>CCGCC--GAG | 3.75±0.59 | 3.89±0.62 | -0.14±0.86 |
| CCAAGΨCUCG<br>GGUU--GGAGC | 4.26±0.55 | 3.89±0.62 | 0.37±0.83 |
| GUCCGΨCUC<br>CAGG--GGAG | 4.94±0.54 | 3.89±0.62 | 1.05±0.82 |
| GUCAGΨCUC<br>CAG--CGGAG | 4.28±0.53 | 3.89±0.62 | 0.39±0.82 |

[a]This loop was excluded from analysis because the loop is much more stable than all other bulge loops.

**Table S12.** Internal loop stabilities. The internal loop is shown in red, and the closing base pairs are shown in blue. All values are shown in kcal/mol.

| Sequences (5'-3') | $\Delta G°_{37\ loop}$ Measured[a] | $\Delta G°_{37\ loop}$ (U analog) | $\Delta G°_{37\ loop}$ Predicted[b] | $\Delta\Delta G°_{37\ loop}$ (Measured – Predicted) |
|---|---|---|---|---|
| UCAGΨCAGU<br>AGUCΨGUCA | -2.74±0.53 | -0.7±0.41 | -4.06±0.81 | 1.32±0.97 |
| GCAΨUCG<br>CGUUAGC | -0.67±0.37 | 0.38±1.32 | -1.61±1.48 | 0.94±1.52 |
| CGCΨGCG<br>GCGΨCGC | -1.93±0.48 | 0.42±0.40 | -2.94±0.80 | 1.01±0.94 |
| GCAΨUUGC<br>CGUUΨACG | -2.14±0.44 | 0.6±0.12 | -1.39±0.69 | -0.75±0.81 |
| GCAΨΨUGC<br>CGUΨΨACG | -3.47±0.48 | 0.6±0.12 | -2.76±0.71 | -0.71±0.86 |
| GCGUΨUCGC<br>CGCUΨUGCG | -0.88±0.45 | 0.83±0.42 | 0.9±1.04 | -1.78±1.14 |
| GCGΨUUCGC<br>CGCUUΨGCG | -2.67±0.51 | 0.83±0.42 | -1.09±1.24 | -1.58±1.34 |
| GCGΨUΨCGC<br>CGCΨUΨGCG | -4.15±0.55 | 0.83±0.42 | -2.46±1.25 | -1.69±1.37 |
| GCAUUΨGC<br>CGΨUUACG | -0.55±0.76 | 0.6±0.12 | -0.36±0.23 | -0.20±0.79 |
| GCAUΨΨGC<br>CGΨΨUACG | -2.15±0.78 | 0.6±0.12 | -0.36±0.23 | -1.80±0.82 |
| UCAGΨCAGU[c]<br>AGUCUGUCA | -1.37± 0.49 | -0.7 ± 0.41 | -2.69 ± 0.79 | 1.32±0.93 |
| UCAGΨCAGU[c]<br>AGUCCGUCA | -0.36 ± 0.46 | 0.32± 0.40 | 0.32 ± 0.40 | -0.68±0.61 |
| CΨGCΨGG[d]<br>GGΨCGΨC | -3.9±0.92 | 0.94± 1.01 | 0.94±1.02 | -4.84±1.38 |

[a]Determined from the experiments.

[b]Determined from the terms for this model.

[c]Reported previously [67].

[d]This sequence was excluded from further analysis because the internal loop appeared to be too stable in relation to other 1×1 nucleotide internal loops.

**Table S13:** Hairpin loop optical melting experiments results. ΔH° is the folding enthalpy change, ΔS° is the folding entropy change, and $\Delta G°_{37}$ is the folding free energy change at 37 °C, determined from the enthalpy and entropy changes. $T_M$ is the melting temperature. These duplexes were melted at the Institute of Bioorganic Chemistry of the Polish Academy of Sciences. Pseudouridines are shown in red.

| Sequences | -ΔH° (kcal/mol) | -ΔS° (eu) | $-\Delta G°_{37}$ (kcal/mol) | $T_M$ (°C) |
|---|---|---|---|---|
| CGUGUUCGAUCCACG | 28.1±5.2 | 85.4±15.6 | 1.56±0.38 | 55.3 |
| CGUGUΨCGAUCCACG | 36.3±4.1 | 110.7±12.4 | 2.01±0.27 | 55.1 |
| CAGACUGAAGAUCUG | 16.1±3.5 | 51.0±10.7 | 0.24±0.13 | 41.8 |
| CAGACUGAAGAΨCUG | 27.8±6.6 | 82.6±19.7 | 2.15±0.53 | 63.0 |
| GGAUUAAUUUCC | 40.7±2.0 | 126.3±6.2 | 1.58±0.14 | 49.5 |
| GGAUUAAUUΨCC | 42.5±3.5 | 130.6±10.6 | 1.95±0.25 | 51.9 |
| GGAΨUAAUUUCC | 45.4±3.8 | 138.1±11.7 | 2.53±0.19 | 55.3 |
| GGAΨUAAUUΨCC | 43.1±5.5 | 129.8±16.9 | 2.89±0.28 | 59.3 |
| GGACUUCGGUCC | 42.2±5.8 | 120.7±16.7 | 4.78±0.61 | 76.6 |
| GGACΨUCGGUCC | 67.3±1.6 | 191.2±4.6 | 8.05±0.18 | 79.1 |
| GGACUΨCGGUCC | 44.5±3.5 | 129.5±10.4 | 4.43±0.34 | 71.5 |
| GGACUUCGGΨCC | 50.7±2.1 | 144.0±6.1 | 5.99±0.20 | 78.6 |

**Table S14.** Hairpin loop stabilities. Pseudouridines are shown in red.

| Sequence | $\Delta G°_{37}$ Measured (kcal/mol) | $\Delta G°_{37}$ Estimated (kcal/mol) | $\Delta\Delta G°_{37}$ (Measured – Estimated) (kcal/mol) |
|---|---|---|---|
| CGUGUUCGAUCCACG | 5.15±0.14 | 5.21±0.64 | -0.06±0.66 |
| CGUGUΨCGAUCCACG | 4.70±0.15 | 5.21±0.64 | -0.51±0.66 |
| CAGACUGAAGAUCUG | 5.85±0.11 | 5.57±0.74 | 0.28±0.75 |
| CAGACUGAAGAΨCUG | 4.06±0.28 | 4.79±1.05 | -0.73±1.09 |
| GGAUUAAUUUCC | 3.58±0.11 | 3.73±0.98 | -0.15±0.99 |
| GGAUUAAUUΨCC | 3.33±0.27 | 3.18±1.03 | 0.16±1.06 |
| GGAΨUAAUUUCC | 2.63±0.14 | 2.67±1.57 | -0.04±1.58 |
| GGAΨUAAUUΨCC | 2.39±0.28 | 3.22±1.18 | -0.83±1.22 |
| GGACUUCGGUCC | 3.07±0.22 | 2.21±0.37 | 0.86±0.43 |
| GGACΨUCGGUCC | -0.20±0.34 | 2.21±0.37 | -2.41±0.50 |
| GGACUΨCGGUCC | 3.42±0.21 | 2.21±0.37 | 1.21±0.42 |
| GGACUUCGGΨCC | 2.81±0.42 | 2.21±0.37 | 0.60±0.56 |

**Table S15:** The helical stack nearest neighbor parameters for Ψ-A pairs compared with parameters that were previously published for single substituted Ψ-A stacks [38]. For each pair, the top sequence is oriented in the 5′ to 3′ direction and the bottom sequence is written 3′ to 5′.

| Stack | $\Delta G°_{37}$ New | $\Delta G°_{37}$ Previous (single substitution) | $\Delta\Delta G°_{37}$ New – Previous (single substitution) |
|---|---|---|---|
| AC<br>ΨG | -3.10±0.23 | -3.29±0.15 | 0.19±0.27 |
| AG<br>ΨC | -2.69±0.23 | -2.77±0.15 | 0.08±0.27 |
| GA<br>CΨ | -2.44±0.25 | -2.49±0.15 | 0.05±0.29 |
| CA<br>GΨ | -2.43±0.25 | -2.20±0.15 | -0.23±0.29 |
| AΨ<br>UA | -2.39±0.33 | -2.80±0.15 | 0.41±0.36 |
| UA<br>AΨ | -2.13±0.29 | -2.10±0.15 | -0.03±0.33 |
| AA<br>UΨ | -2.10±0.33 | -2.74±0.15 | 0.64±0.36 |
| AA<br>ΨU | -1.38±0.28 | -1.62±0.15 | 0.24±0.32 |

**Table S16:** The helical stack nearest neighbor parameters for Ψ-A pairs compared with parameters that were previously published for fully substituted Ψ-A stacks [10]. For each pair, the top sequence is oriented in the 5′ to 3′ direction and the bottom sequence is written 3′ to 5′.

| Stack | $\Delta G°_{37}$ New | $\Delta G°_{37}$ Previous (full substitution) | $\Delta\Delta G°_{37}$ New – Previous (full substitution) |
|---|---|---|---|
| AA<br>ΨΨ | -0.92±0.19 | -1.23±0.05 | 0.31±0.20 |
| AΨ<br>ΨA | -1.94±0.44 | -1.52±0.14 | -0.42±0.46 |
| ΨA<br>AΨ | -1.53±0.42 | -1.71±0.16 | 0.18±0.45 |
| AG<br>ΨC | - 2.69±0.23 | -2.10±0.10 | -0.59±0.25 |
| CA<br>GΨ | -2.43±0.25 | -2.35±0.08 | -0.08±0.26 |
| AC<br>ΨG | -3.10±0.23 | -2.50±0.08 | -0.60±0.24 |
| GA<br>CΨ | -2.44±0.22 | -2.51±0.10 | 0.07±0.27 |
